# Organoid-based *in vitro* system and reporter for the study of *Cryptosporidium parvum* sexual reproduction

**DOI:** 10.1101/2023.09.29.560165

**Authors:** Bethany R. Korwin-Mihavics, Emmett A. Dews, Peter Miller, Alexandra Cameron, Bruno Martorelli di Genova, Christopher D. Huston

## Abstract

Many advances have been made recently in our understanding of *Cryptosporidium*’s asexual cycle and sexual differentiation. However, the process of fertilization, which is required for transmission of infectious oocysts, is not well understood. Typical cancer cell-based culture only allows robust exploration of asexual cycle and sexual differentiation of *Cryptosporidium*. To facilitate exploration of sexual reproduction in *C. parvum* we developed an organoid-based culture system that supports *Cryptosporidium’s* full life cycle and a novel fertilization reporter. Organoid derived monolayers (ODMs) supported fertilization and oocyst production and maintained the infection for up to 3 weeks. ODM derived oocysts were infectious *in vivo*. Fertilization was confirmed by successfully mating two strains of *C. parvum* and with a novel fertilization switch reporter. The fertilization switch reporter utilizes a DiCre system in which cre fragments are expressed under the control of sexual stage promoters resulting in a rapamycin-inducible switch in fluorescent protein expression from mCherry to mNeonGreen after fertilization that is spatially and temporally controlled. This results in mCherry positive parasites in the first generation and offspring that express mNeonGreen. *In vivo* validation of the fertilization switch reporter demonstrated the precision and efficiency of the fertilization switch reporter and confirmed excision of the mCherry gene sequence only after rapamycin treatment. The start of a second generation of parasites was also shown in the ODMs and rarely in HCT8s. Use of this reporter in ODMs can help investigate the *Cryptosporidium* lifecycle post sexual differentiation in a physiologically relevant *in vitro* system.

**Importance:** Organoid derived monolayers provide an opportunity to elucidate previously inaccessible aspects of *Cryptosporidium*’s biology. This system overcomes the disadvantages of previous organoid-based methods for *Cryptosporidium* culture. It is faster and simpler than previously described systems, uses defined media to increase reproducibility and consistency, enables real-time observation, supports parasite fertilization and oocyst production, and provides a physiologically relevant tissue culture system to facilitate studies of *Cryptosporidium* cell biology. The ODM system could facilitate the study of host-pathogen interactions, *Cryptosporidium*-host specificity, or innate or cellular immune responses to *Cryptosporidium* infection stimulated in the intestinal epithelium. The fertilization switch reporter could be used to test factors or drugs that may have potential to interfere with *Cryptosporidium’s* sexual reproduction. Organoid-based cell cultures in combination with the fertilization switch reporter could increase our understanding of sexual reproduction in *Cryptosporidium,* leading to vital information for the development of sexual reproduction inhibitors or vaccines that could shorten disease duration and prevent transmission.

## Introduction

*Cryptosporidium* is an intestinal protozoan parasite that is a major cause of diarrheal disease worldwide and is among the 5 most common infectious causes of moderate to severe diarrhea in children under 2 years^1–4^. Although not officially designated a neglected tropical disease, the morbidity and mortality of *Cryptosporidium* outpaces that of many officially recognized neglected tropical diseases^5^. Half of all waterborne outbreaks in the United States are caused by *Cryptosporidium*, with *C. hominis* and *C. parvum* being the main species that cause outbreaks^6^. *Cryptosporidium* is found on every continent but Antarctica, but the heaviest burden of disease falls on low-income nations and low-resource communities, especially in sub-Saharan Africa^1,3,5^. In immunocompetent adults, cryptosporidiosis is self-limited and usually resolves within 2-3 weeks, but immunocompromised individuals and young children, particularly malnourished young children, are the most prone to developing severe disease that can become chronic and/or life-threatening^1,2,5^. Currently, there is no vaccine and nitazoxanide, the only drug approved for cryptosporidiosis, is poorly effective in the most vulnerable populations prone to severe cryptosporidiosis. Chronic or repeated infection with *Cryptosporidium* can cause intestinal damage resulting in loss of absorptive surface area and contribute to future malabsorption and malnutrition, resulting in a vicious cycle of malnutrition and infection that can result in long-term growth stunting and cognitive deficits in young children^7^.

*Cryptosporidium* exhibits monoxenous development, whereby the entire lifecycle is completed within a single host. Upon ingestion, oocysts pass through the acidic stomach environment and into the small intestine where bile acids and digestive enzymes promote excystation to release motile sporozoites^8,9^. Sporozoites invade the host cell, establish an epicellular niche in which the parasite is intracellular but extracytoplasmic, and develop into trophozoites that mature into meronts containing 8 merozoites which egress and reinvade new host cells^9–11^. After three rounds of merogony, merozoites differentiate into sexual stages, the microgamont (male stage) and the macrogamont (female stage)^10,12^. Microgamonts release microgametes which find and fertilize a macrogamont forming a zygote which sporulates to form a new oocyst. Oocysts may excyst within the gut lumen and begin a new round of infection or be excreted into the environment where they wait to be ingested by a new host^10–12^.

Due to its monoxenous development, a system of continuous culture should be simple, but such a system has been elusive for *Cryptosporidium*. Approaches that have supported oocyst production have not been widely adopted, suggesting excessive complexity, incompatibility with study needs, or an inability to reproduce the results. Conventional HCT8 culture supports asexual reproduction and sexual differentiation, but robust production of new oocysts is not supported, suggesting a block to fertilization and zygote formation^7,12^. Culturing HCT8s in a candle jar has been reported to support the production of thin-walled oocysts^13^. Culture in a bioreactor with hollow nanofibers lined with HCT8s produced infectious oocysts for up to 20 weeks and maintained oocyst production for 2 years, but this has not been widely adopted and is not compatible with microscopy methods^14,15^.

Organoid-based infection models offer a more promising and physiologically relevant approach due to the presence of differentiated intestinal epithelial stem cells. One study using three dimensional organoids derived from human intestinal epithelial cells showed new oocyst production but required tedious microinjection of the enteroids for the infection^16^. Another intestinal stem cell derived system utilized targeted laser ablation to create a Matrigel™ channel that mimicked intestinal crypt and villi structures to serve as a scaffold for intestinal stem cell attachment and differentiation and reported oocyst production for 4 weeks^17^. A stem cell-derived air-liquid interface culture system also reported new oocyst production but required the maintenance of two cell lines and the initial cell differentiation took up to 11 days^18^. The method described here has been adapted from a number of protocols to support the full lifecycle of *Cryptosporidium in vitro*^19–22^. Importantly, our method uses a defined media with readily available reagents and equipment to support complete *Cryptosporidium* lifecycle development physiologically relevant intestinal environment.

CRISPR/Cas9 systems enabling genetic modification^23,24^, RNA-Seq and single cell RNA sequencing data sets refining our knowledge of sexual differentiation^10,12,25,26^ and new culture systems have opened the possibility of studying fertilization *in vitro*^16,18^. The programmed nature of *Cryptosporidium*’s life cycle in which sexual differentiation occurs after completion of three rounds of asexual reproduction means that sexual reproduction is required for continued expansion within the host, which makes treatments or vaccines targeting fertilization mechanisms extremely attractive as they could shorten the disease and also help prevent transmission^10^.

Past approaches to measuring fertilization have relied on mating of different strains of *C. parvum*^12,18^. While excellent approaches, they only capture interstrain crossing and we sought to develop a reporter that would detect all fertilization events, including fertilization within the same parasite strain. Here we present a novel fertilization reporter for use in studying *Cryptosporidium* fertilization.

## Results

### Organoid derived monolayers exhibit the characteristics of differentiated intestinal epithelium and support *C. parvum* infection

Confluent mouse intestinal stem cells were grown on Matrigel-coated permeable supports with 50% stem cell maintenance media (SCMM) for the first 24 hours, 5% SCMM for the second 24 hours, and were then allowed to differentiate in differentiation media (DM) for 2 to 3 days before *C. parvum infection* (Fig. 1A). Characteristics of functional intestinal epithelial tissue, e.g., microvilli (Fig. 1J and 1K) and tight junctions (Fig. 1B, F, and K), and presence of differentiated cell types were confirmed by immunofluorescence microscopy and gene expression. Goblet, enteroendocrine, and Paneth cells were identified by staining for Muc2, Chromogranin A, and Lysozyme, respectively (Fig 1 G-I). ZO-1 gene expression, indicating tight junction formation, was unchanged between spheroids and ODMs (Fig. 1B). Stem cell marker Lgr5 gene expression was significantly reduced in differentiated ODMs compared to spheroids. *Muc2* gene expression was significantly increased in ODMs compared to spheroid culture, and chromogranin A (*ChgA*) expression was elevated in ODMs, although not significantly (Fig. 1C-D). Altogether, these data confirmed differentiation of the ODMs after 2-3 days in DM. Turnover of the ODMs was also observed. During media changes, dead cells were observed in spent media, but the monolayers remained intact, indicating that dead cells that had sloughed off were likely replaced with new cells (Supplemental Fig. 1D). The cell turnover rate appeared comparable to that of intestinal epithelium *in vivo* (2-3 days in mice and 3-5 days in humans)^27^.

**Figure 1:**
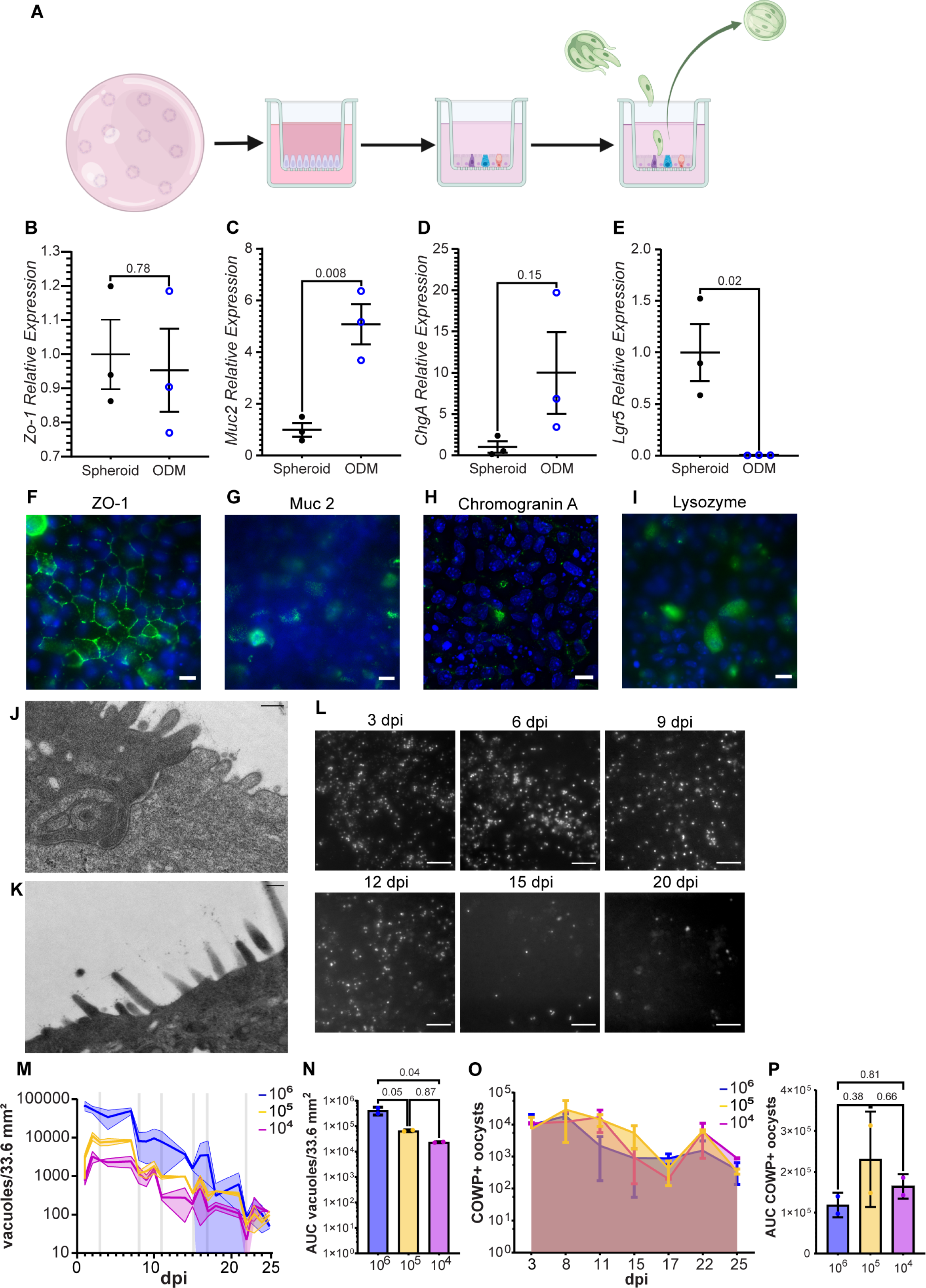
Organoid derived monolayers exhibit intestinal epithelium characteristics and support *C. parvum* infection. (**A**) Schematic for organoid derived monolayer (ODM) differentiation and infection. Mouse intestinal stem cells (mISC) were embedded and grown in Matrigel domes and maintained as spheroids in SCMM until expanded enough to seed permeable supports. Prior to seeding, permeable supports were coated with a 1:10 Matrigel in SCMM and incubated overnight at 4°C. Matrigel dilution was removed and permeable supports were incubated at 37°C for at least 15 minutes prior to seeding with mISC. Permeable supports were seeded with 3x10^5^ cells/well. Once confluent, the SCMM concentration was reduced to 5% for 24 hours, and then differentiation media (DM) which lacked L-WRN conditioned media entirely. ODMs were maintained in DM for 2-3 days and then infected with *C. parvum*. (**B-E**) qRT-PCR of mRNA expression of ZO-1 for tight junctions (**B**), Muc2 for goblet cells (**C**), ChgA for enteroendocrine cells (**D**), and lgr5 for stem cells (**E**). A Student’s t-test was used to assess statistical significance with a cut-off of 0.05. P-values are displayed on each graph. (**F-I**) Immunofluorescent staining of ZO-1 (**F**), Muc2 (**G**), Chromogranin A (**H**), and Lysozyme (**I**) in green and nuclei in blue confirming the presence of tight junctions, goblet cells, enteroendocrine cells, and Paneth cells respectively in differentiated ODMs. Scale bars are 10 µm. (**J-K**) TEM confirmation of tight junction (**J**) and microvilli (**K**) on ODMs. Scale bars are 200 nm. (**L**) Representative live epifluorescence images of ODM infection with mNeonGreen parasites 3, 6, 9, 12, 15, and 20 dpi. Scale bar is 50 µm. (**M**) Vacuole counts from image J macro analysis of fluorescent parasite vacuoles adjusted for total cell culture area. Lines represent infection with different infectious doses (10^6^ oocysts (blue), 10^5^ oocysts (yellow) and 10^4^ oocysts (magenta). Vertical grey lines show days of supernatant collection (3, 8, 11, 15, 17, 22, and 25 dpi); images were taken after supernatants were collected. Standard deviation is shown as shaded area surrounding each line. (**N**) Area under the curve (AUC) of experiment in M. (**O**) Flow cytometry quantification of COWP1 + oocysts collected from ODM supernatant. (**P**) AUC of oocyst counts in O. One-way ANOVA (Tukey post-hoc multiple comparisons) was used to assess statistical significance of AUC in N and P with a p-value cut-off of 0.05. P-values are displayed on graphs. Error bars show standard deviation. n=2 for each condition.

Differentiated monolayers infected with 10^6^, 10^5^, or 10^4^ mNeonGreen expressing *C. parvum* oocysts were tracked over the course of 3 weeks with live widefield epifluorescence microscopy (representative images in Fig. 1L). The parasite vacuole (PV) load decreased slowly over the course of 3 weeks (Fig. 1M). By 21 dpi, PVs were still present, but they were no longer present in each field of view. The biggest drops in vacuole load were concurrent with supernatant collections (Fig. 1M), and after an initial drop, the population often increased and leveled out until the next supernatant collection was performed. These trends occurred in multiple similar experiments (Supplementary Fig. 1). Over the course of the infection, an average of about 4.1 x10^5^, 6.6 x10^4^, and 2.3 x 10^4^ PVs were estimated to have formed per monolayer based on calculating the area under the curve of PVs for the total infection course for 10^6^, 10^5^, or 10^4^ infective doses, respectively, with the estimated total PVs for the 10^6^ dose being significantly higher than the others (Fig. 1N). Oocysts collected from the supernatant of ODMs were quantified by flow cytometry (Fig. 1O). Even though the 10^6^ infective dose resulted in a significantly higher number of estimated PVs (Fig. 1M-N), this did not correlate with the number of oocysts produced (Fig. 1O-P) compared to the smaller infective doses. In fact, oocyst output for each condition was comparable (Fig. 1P). Oocyst counts did not directly increase with the drop in number of vacuoles observed after supernatant collection, likely because many of the host cells that sloughed off the monolayer were also infected (Fig. 1D). Overall, organoid derived monolayers support robust infection with *C. parvum* and production of new oocysts.

### ODM derived oocysts are infectious *in vivo*

To verify the infectiousness of oocysts derived from ODMs, HCT8 and ODM cultures were infected with tdTomato-nLuc *C. parvum*. The parasites were excysted and filtered to exclude any unexcysted oocysts and ensure only free sporozoites were used to infect the cultures and any oocysts collected from the cultures were newly generated. Approximately equivalent surface areas were used for each set of cultures. Supernatants were collected for 2 weeks, then concentrated by centrifugation and used to gavage 5 IFNγ-/- mice for each condition (Fig. 2A). Mouse fecal collections were performed over the course of 3 weeks. Nanoluciferase assay analysis of the fecal samples indicated that all 5 mice gavaged with ODM derived supernatants were infected, while none of the mice gavaged with HCT8 derived supernatants were ever positive for nanoluciferase in the feces (Fig. 2B). This indicates that ODMs support the *in vitro* development of new infectious oocysts in great enough numbers to support subsequent *in vivo* infections.

**Figure 2:**
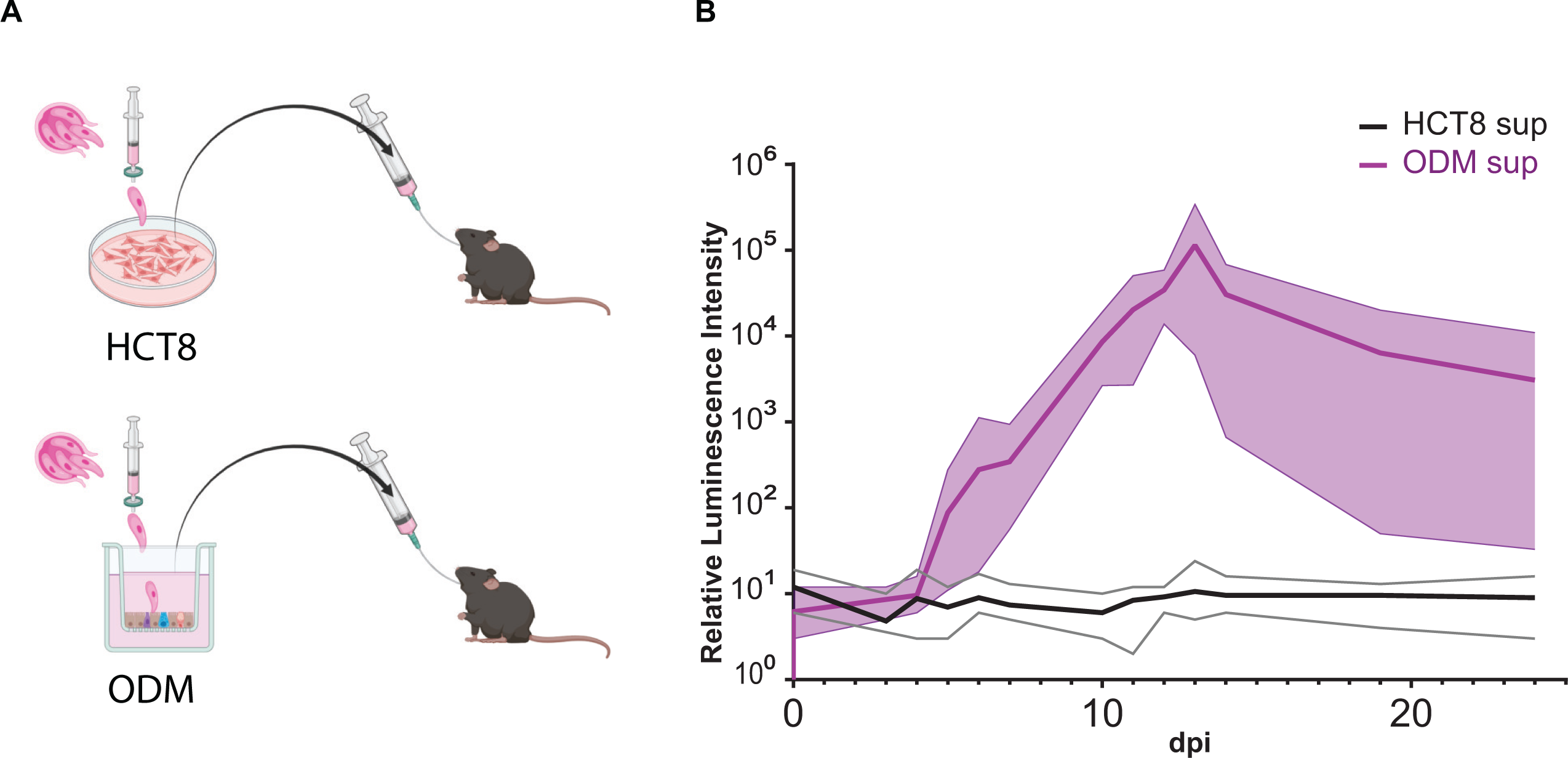
Organoid derived monolayers support *C. parvum* production of infectious oocysts. **(A)** Graphical experimental representation. tdTomato-nLuc-Neo oocysts were excysted and free sporozoites were purified by filtration through a 1.2 µm pore syringe filter. Purified sporozoites were used to infect 24 wells (∼33 mm^2^ each) of typical HCT8 culture or ODMs (1.5 x 10^5^ oocysts/well). Supernatants were collected every other day for 2 weeks, pooled and concentrated by centrifugation, and stored at 4°C. IFNγ -/- mice were gavaged with supernatants from either HCT8 culture or ODM culture. Fecal samples were collected on days 1-14, 19 and 24 post infection and fecal nanoluciferase was assessed. **(B)** Nanoluciferase assay of fecal samples collected from mice gavaged with infected HCT8 or ODM supernatants. Thick lines represent the mean Relative Luminescence Units (RLU) for each group and thin lines represent the range of RLU for each condition. Results are representative of 1 experiment, n=5 mice per group.

### Organoid derived monolayers support *C. parvum* mating

Organoid derived monolayers were coinfected with mNeonGreen (mNG) and tdTomato (tdT) reporter *C. parvum* strains and infections were observed and recorded with live imaging over the course of 3 weeks. Double positive vacuoles indicative of mating were present (Fig. 3A), but rare compared to single positive vacuoles (Fig. 4A-C). Oocyst output from ODMs can serve as a proxy for fertilization and total oocyst output from the 3 experiments was not statistically significantly different (Fig. 4G-H). New oocysts with single positive, double positive, and double negative phenotypes were also produced from these infections (Fig. 4D-F). Double positive oocysts were again rare events and were not observed in all supernatant collections (Fig. 4D-F, Fig 4B). While most of the oocyst population collected in supernatant from the ODMs was composed of double negative oocysts, we cannot confirm that they resulted from outcrossing because a high proportion of double negative oocysts were detected in the washes performed immediately after infection (Sup Fig. 3B), suggesting that they were present in the input. Upon retrospective examination of the tdT and mNG stocks used for these experiments, the mNG stock was confirmed to be a mixed population of mNG and non-fluorescent oocysts. This means that in truth the results reflect the mixing of 3 phenotypes rather than 2 and that vacuole counts from epifluorescence microscopy lacked data for the double negative population and underrepresented the total parasite load. Regardless of the limitations, the presence of double positive vacuoles within the ODM and double positive oocysts in the supernatant combined with the total oocyst output and infection of mice with ODM-derived oocysts confirm that fertilization and oocyst production is supported in organoid derived monolayers.

**Figure 3:**
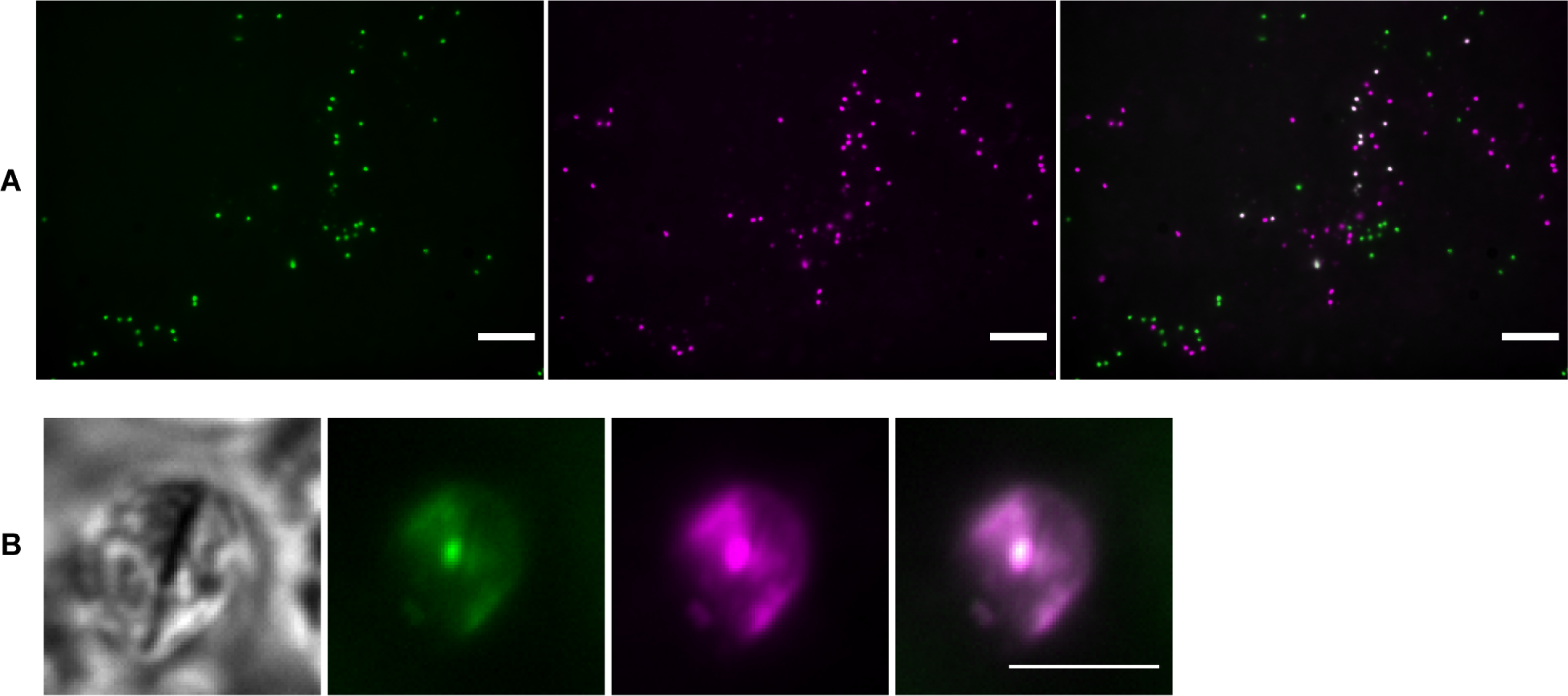
Interstrain mating confirmed by double positive zygote and oocyst observation. (**A**) Live image of ODM infected with mNeonGreen (mNG) (left) and tdTomato (tdT) (middle) expressing *C. parvum* strains 8 dpi. Merge on the right shows double positive vacuoles in white. Scale bar is 50 µm. Image was processed for 2D deconvolution using AutoQuant X. (**B**) Oocyst collected from supernatant expressing both mNG and tdT. Scale bar is 5 µm.

**Figure 4:**
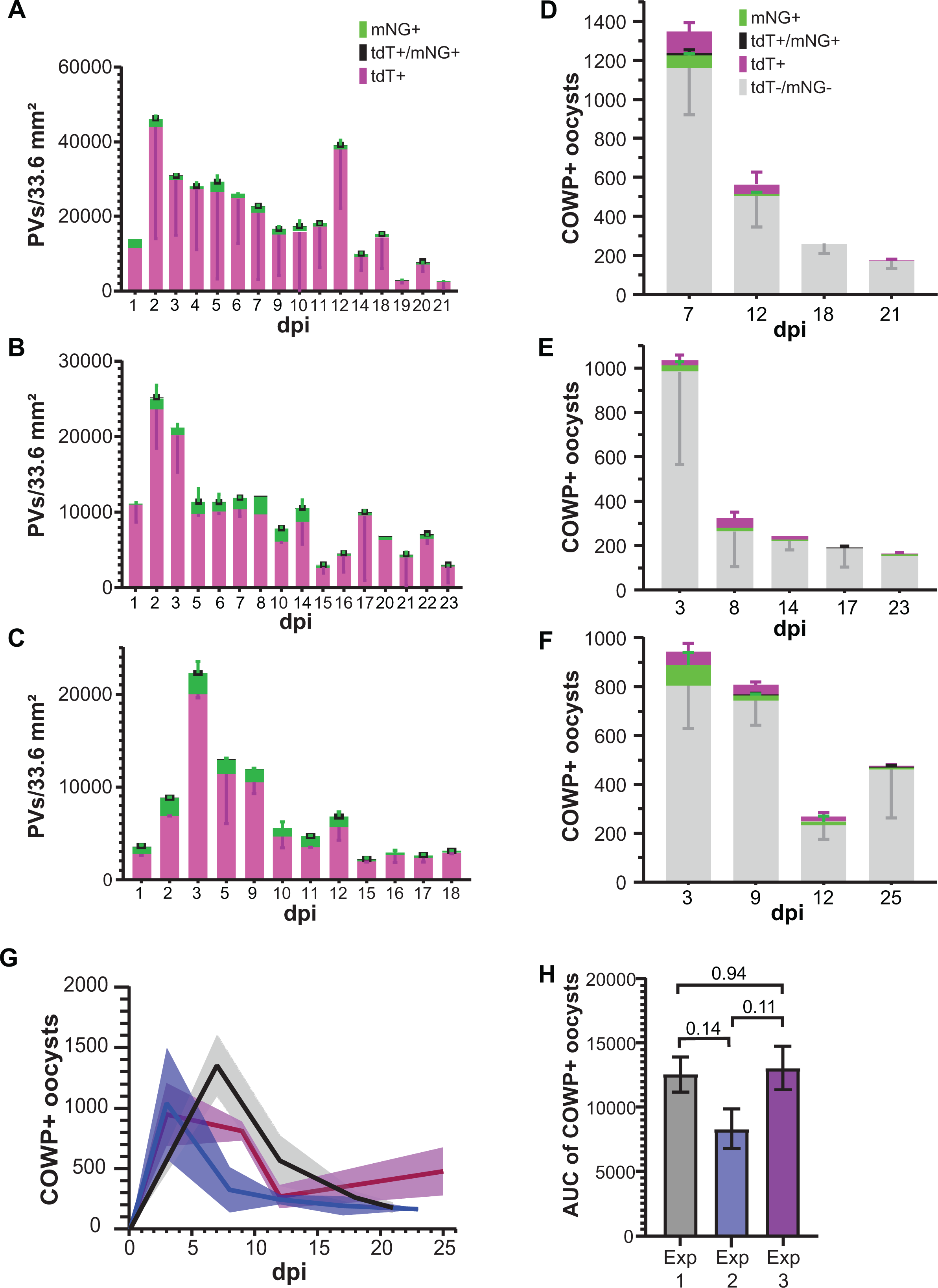
Successful mating of two transgenic *C. parvum* strains in organoid derived monolayers. (**A-C**) Quantification of parasitophorous vacuoles (PVs) from Image J macro adjusted for total cell culture area. Each graph shows results from a different experiment. (**D-F**) Flow cytometry quantification of total oocysts collected from supernatant. Each graph shows oocyst results from individual experiments in A-C respectively. Error bars show standard deviation. (**G**) Total COWP1+ oocysts from supernatant over the course of the experiment. Each line represents a different experiment. Standard deviation is shown as shaded area surrounding each line. (**H**) AUC of COWP1+ oocysts for each experiment shown. One-way ANOVA (Tukey post-hoc multiple comparisons) used to assess statistical significance. P-values are displayed on graph. P-values ≤ 0.05 were considered significant. Error bars show standard deviation. n=2 for each experiment. Data is representative of 2 experiments.

### A novel fertilization reporter: Fertilization Switch Reporter

Previous studies have focused on mating 2 strains of parasites to measure fertilization and explore recombination^12,28^. These studies successfully captured events resulting from interstrain crosses, but did not capture intrastrain mating. Similarly, we initially crossed tdTomato and mNeonGreen reporter parasite strains to visualize fertilization (Figures 3-4), which confirmed fertilization but quantification was confounded by presence of non-fluorescent parasites and inability to identify intrastrain mating. To be able to measure all fertilization events including “selfing”, i.e. fertilization within the same parasite strain, we developed a reporter that exhibits a color switch from red to green after fertilization occurs (Figure 5). To this end, we utilized the DiCre system that has been used by others in the parasitology field and has recently been demonstrated to be tractable in *Cryptosporidium*^26,29–31^. The fertilization switch reporter is inducible and spatially and temporally controlled. Cre subunits fused to FKB12 and FRB subunits dimerize in the presence of rapamycin, bringing the Cre subunits together to make a functional CRE recombinase. To gain spatial control of dimerization and precisely measure fertilization, one Cre subunit was expressed in the microgametocyte under control of the HAP2 promoter (cgd8_2220) and the other expressed only in the macrogamont under control of the COWP1 promoter (cgd6_2090). In this way, the Cre subunits are only expressed in the sexual stages and are only in the same physical space after fertilization and during zygote development. In the presence of rapamycin, functional Cre recombinase will assemble in zygotes and excise a floxed mCherry-STOP sequence under control of a constitutive aldolase promoter (cgd1_3020), leaving behind an mNeonGreen sequence. A switch from red to green parasites in the second generation should be observed.

**Figure 5:**
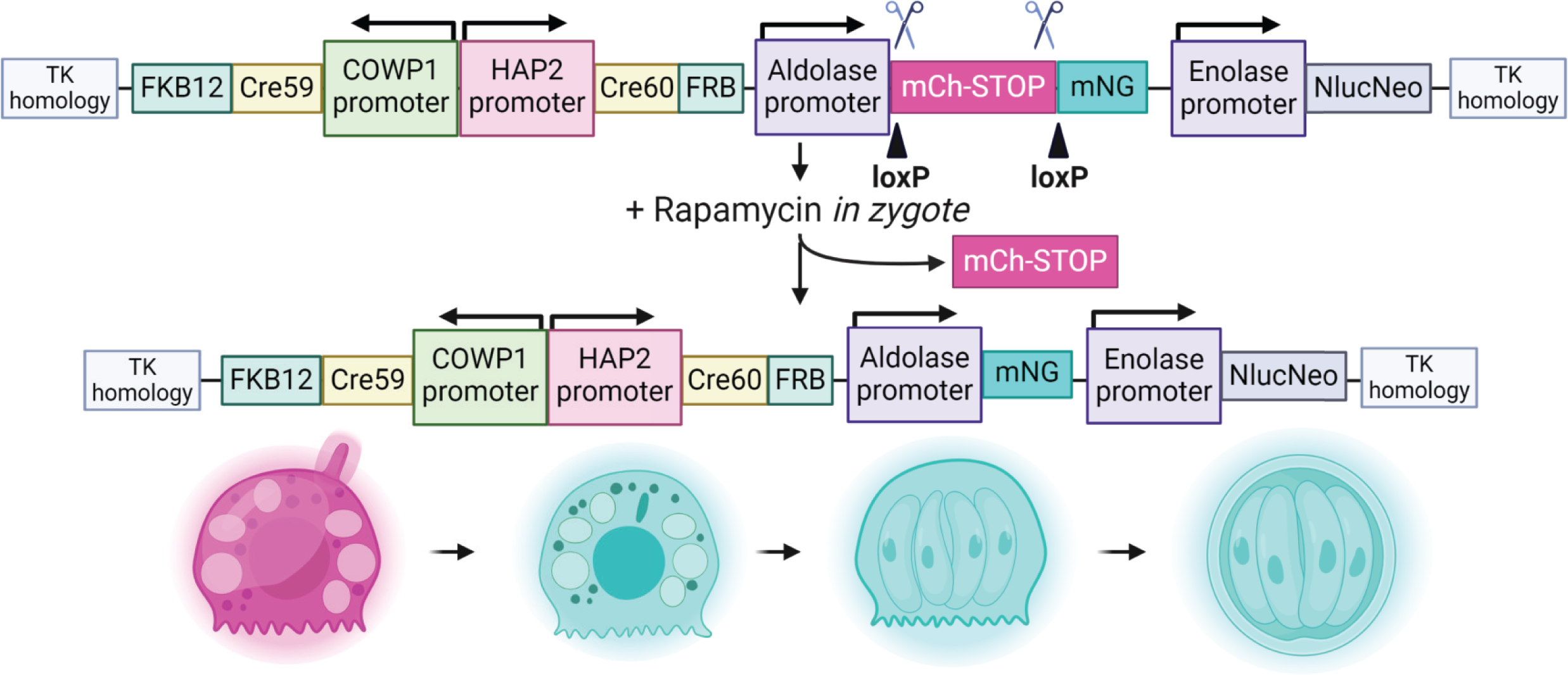
Fertilization Switch reporter. Sexual stage specific promoters drive the expression of Cre recombinase fragments fused to FRB and FKB12 which will dimerize in the presence of rapamycin. Because they will be spatially separated due to expression in different sexual stages, they cannot dimerize in the presence of rapamycin unless they are in the same cell which occurs only after fertilization. LoxP sites flanking an mCherry sequence immediately upstream of an mNeonGreen sequence will be excised in the presence of rapamycin and when the Cre fragments are in the same physical space in a zygote.

We first sought to validate the functionality of the fertilization switch reporter *in vivo*. Mice infected with the fertilization switch reporter strain were either left untreated or gavaged with 10 mg/kg rapamycin for 4 days (three doses/day for a total of 10 mg/kg) (Fig. 6A). Seven days post treatment, fecal samples were collected, purified, and analyzed. Oocysts in mouse fecal samples were analyzed by epifluorescence microscopy (Fig. 6B) and flow cytometric analysis (Fig. 6C-D). Microscopy confirmed mCherry fluorescence and mNeonGreen fluorescence of oocysts that were overwhelmingly mNeonGreen positive and negative for mCherry following rapamycin treatment (Fig. 6B). An mCherry positive oocyst was observed only once under the microscope in oocyst preparations made from the rapamycin treated group, while oocysts from untreated mouse fecal samples were exclusively mCherry positive. When quantified by flow cytometry, the oocyst population in feces from mice that received rapamycin treatment shifted from mCherry positive to over 95% mNeonGreen positive, 11.5% of which were positive for both mCherry and mNeonGreen (Fig. 6C-D). PCR of the region containing the loxP-mCherrySTOP-loxP-mNeonGreen sequence from purified fecal oocysts showed a band shift consistent with successful excision of mCherry in the rapamycin treated group (Fig. 6E). No unexcised band was detected in the rapamycin treated group. Collectively, the data indicated that the color switch post rapamycin treatment was a highly specific and efficient marker of fertilization.

**Figure 6:**
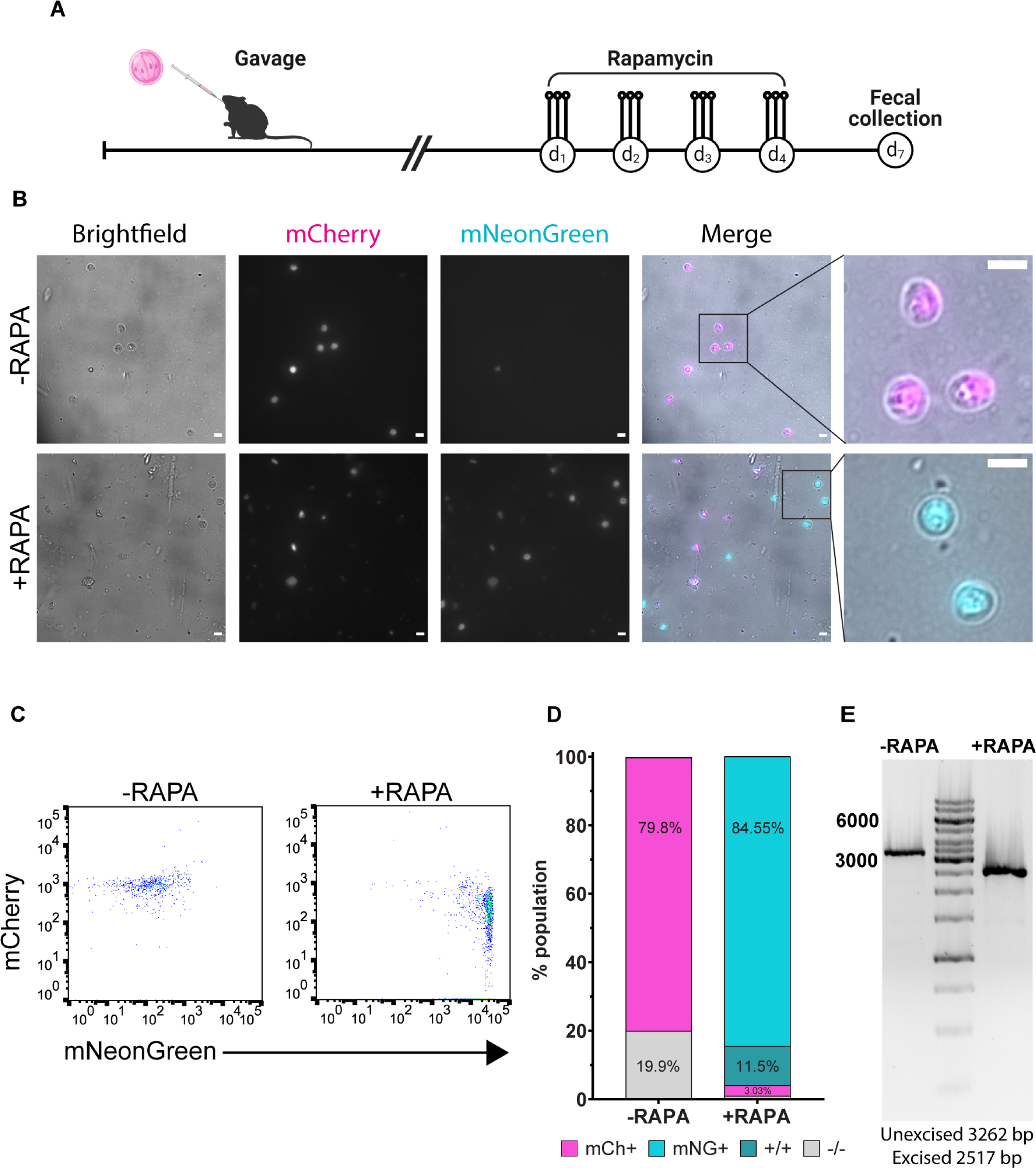
In vivo validation of the Fertilization Switch reporter demonstrates its inducibility and efficiency. (**A**) Oocysts were purified from fecal samples collected from Fertilization Switch reporter infected NSG mice that were left untreated or were treated with 10 mg/kg rapamycin daily for 4 days. Purified oocysts were used for microscopic analysis (**B**), flow cytometry (**C-D**), and PCR analysis (**E**). (**B**) Top row: no rapamycin treatment; bottom row: with rapamycin treatment. Far right column: zoomed in view of oocysts of merged images. mCherry in magenta, mNeonGreen in cyan. Scale bars are 5 µm. (**C**) Flow cytometry plots of purified oocysts collected from mice with or without rapamycin treatment. (**D**) Percentages of COWP positive oocysts expressing mCherry or mNeonGreen. (COWP+/mCherry or mNeonGreen+) (**E**) DNA gel of DNA collected from oocysts purified from fecal samples from mice with (right lane) or without (left lane) rapamycin treatment.

### Fertilization switch reporter functions in ODMs

Next, we sought to test the fertilization switch reporter in ODM culture. We had previously shown interstrain mating in ODMs and wanted to determine if we could capture intrastrain mating in ODMs as well. Unfortunately, the fluorescence of the fertilization switch reporter was not bright enough to readily observe the fertilization reporter live in culture as it was intended. However, by staining with anti-RFP and anti-mNeonGreen antibodies the fluorescent proteins could be detected. After validating the function of the fertilization switch reporter *in vivo*, we wanted to use it to confirm its function in the ODMs.

ODMs were infected with the fertilization switch reporter, wells were washed 16 hpi, and (apical) media was replaced with differentiation media with or without 100 nM rapamycin. A week after infection, the ODMs were fixed, stained, and imaged. In untreated transwells, mNeonGreen positive vacuoles were not observed while between 25-40% of vacuoles in the rapamycin treated ODMs were mNeonGreen positive (Fig 7A-B). Interestingly, mononucleated and multinucleated mNeonGreen positive vacuoles corresponding to the sizes of trophozoites and meronts were also observed, indicative of autoinfection and a second generation of *Cryptosporidium* in the ODMs (Figure 8).

**Figure 7:**
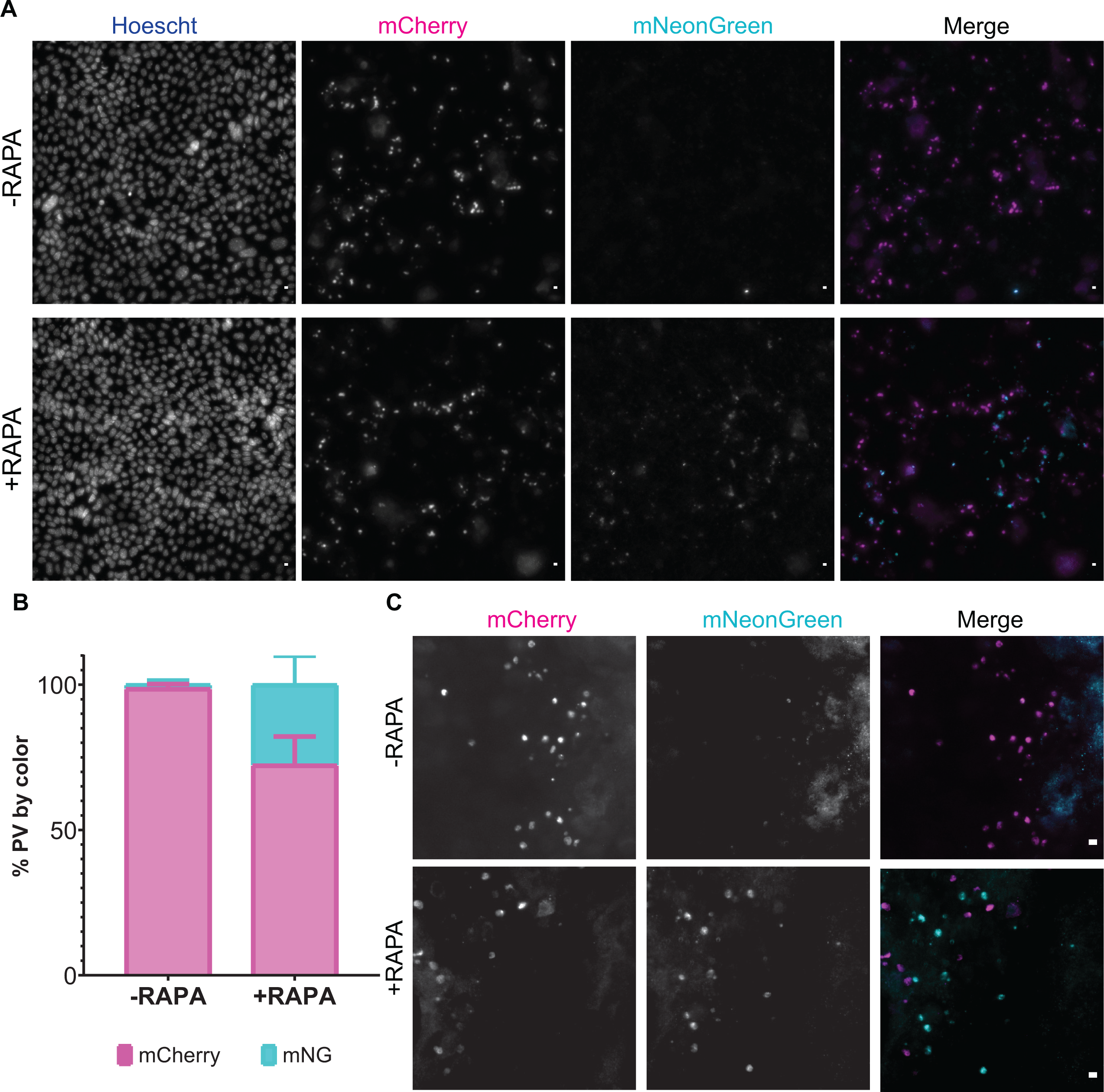
Fertilization switch reporter infection of ODMs. (**A**) Representative images of fertilization switch reporter infected ODMs with (bottom row) or without (top row) rapamycin treatment. (**B**) Percentage of vacuoles of large tiled images counted as mCherry or mNeonGreen positive. Error bars represent SD. (**C**) Higher magnification of infected area showing a mixed population of mCherry and mNeonGreen vacuoles. In merged images, mNeonGreen fluorescence is shown in cyan and mCherry fluorescence is magenta. Data are representative of 2 experiments. Scale bars are 5 µm.

**Figure 8:**
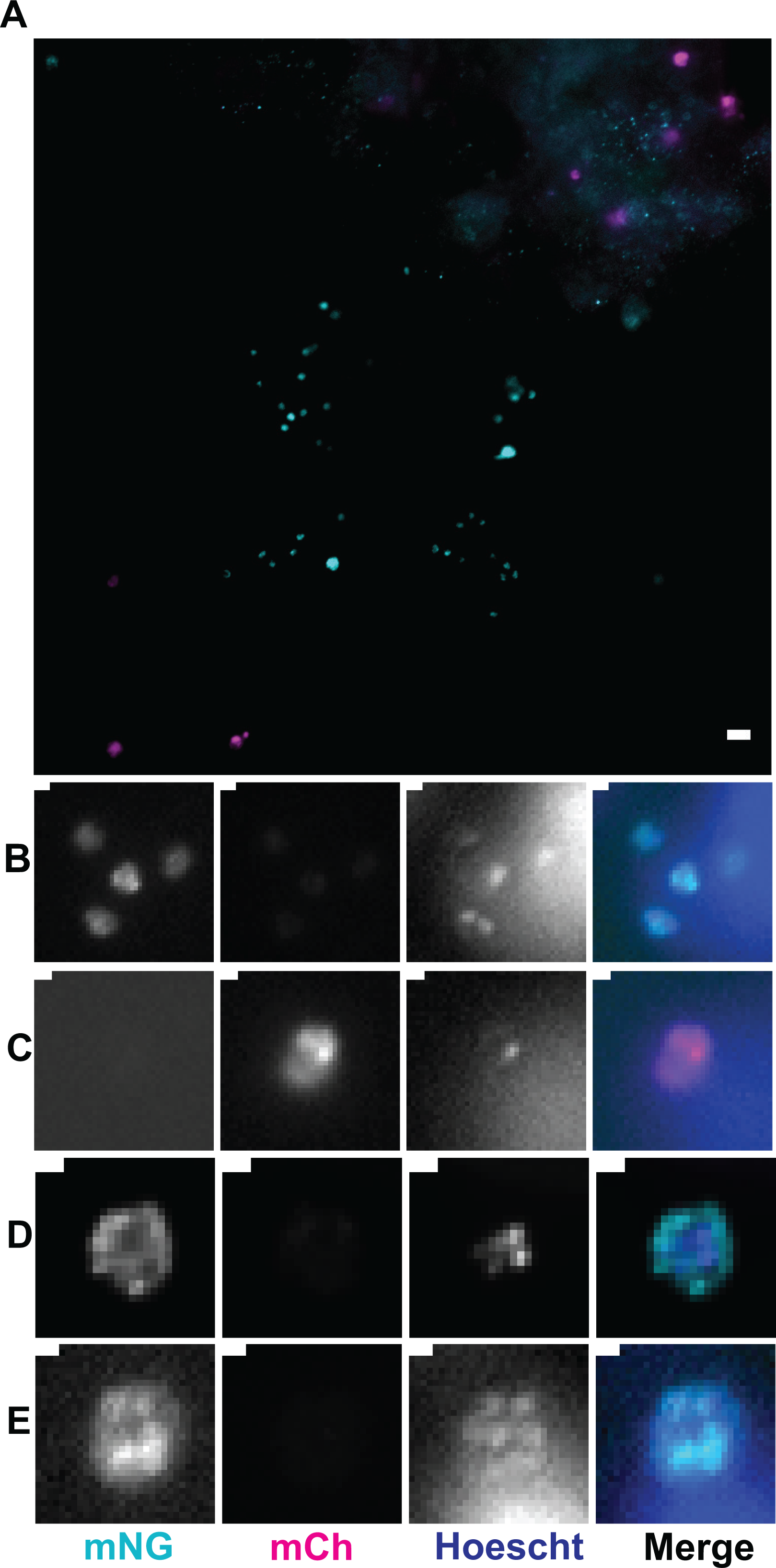
Evidence of autoinfection in ODMs. (**A**) Infected area with a mixed population of mNeonGreen and mCherry positive vacuoles. Scale bar is 5 µm. (**B**) Z-stack slice showing trophozoites. (**C**) Larger mCherry positive vacuole with only 1 nucleus, likely an unfertilized macrogamont. (**D**-**E**) Z-stack slice showing multiple nuclei in a larger mNeonGreen positive vacuole, likely a meront. Scale bars for B-E are 1 µm.

### Fertilization in HCT8 cell monolayers

The fertilization switch reporter was also used to infect HCT8 cells, expecting to confirm a complete block to fertilization. No mNeonGreen clusters were observed in the absence of rapamycin treatment (Fig. 9A). To our surprise, clusters of mNeonGreen positive vacuoles were observed repeatedly following rapamycin treatment, although they were rare compared to mCherry positive vacuoles (Fig. 9B-C; 88 hpi). Based on the number, size, and distribution of vacuoles, these clusters likely represented sites of fertilization followed by a single round of merogony. Interestingly, mNeonGreen positive parasite clusters were not detected at 48 hpi but at 90 hpi were detected more frequently when infecting HCT8 cell monolayers grown in fibronectin-coated MatTek plates (Fig 9D). This could be related to coating the MatTek plates with fibronectin before seeding HCT8s, which may have promoted stronger polarization of the cell monolayer.

**Figure 9:**
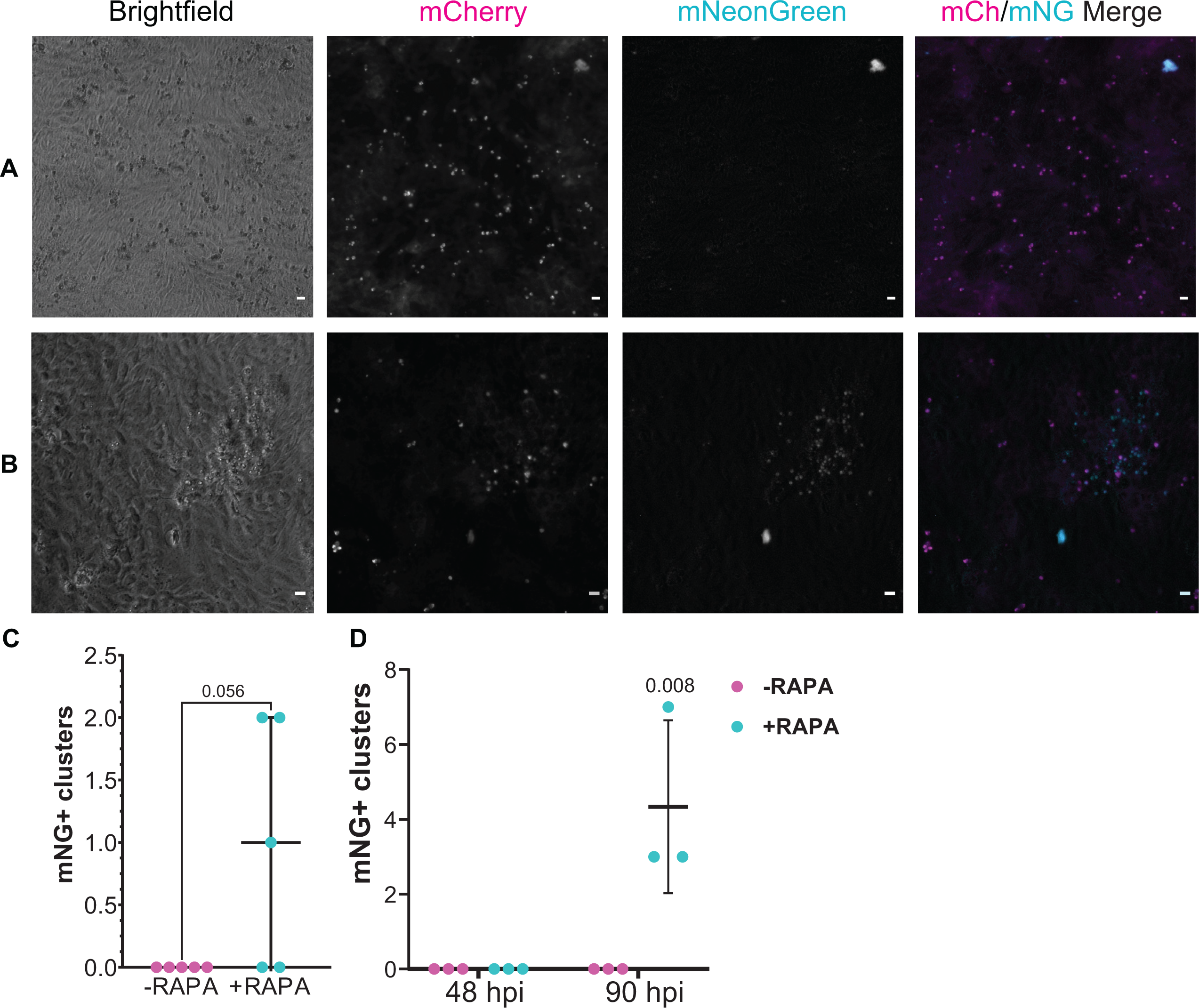
Fertilization switch reporter infection in HCT8 cells. Fertilization switch infected HCT8 cells in the absence (**A**) or presence (**B**) of rapamycin treatment at 88 hpi. (**B**) Representative mNeonGreen positive cluster of parasite vacuoles. Scale bars are 5 µm. (**C**) Number of clusters of mNeonGreen positive parasite vacuoles counted at 88 hpi. Each data point indicates total mNG+ vacuole clusters present on a 33 mm^2^ culture well with data accrued in two independent experiments.

## Discussion

This study shows that organoid-based cultures generated using defined culture media contain differentiated cell types representative of intestinal epithelial tissue and support *Cryptosporidium* infection and sexual reproduction. Organoid derived monolayers (ODMs) support *C. parvum* infection and new infectious oocyst production for at least 21 days and enable studies of fertilization. Important advances specific to this study include use of a defined culture medium that will enable simpler reproduction and use of the system by the community of *Cryptosporidium* researchers, and a precise tool in the fertilization switch reporter parasite with which to quantify fertilization that should enable new insights into the mechanisms underlying the largely unstudied *Cryptosporidium* sexual replication cycle.

Organoid-based models represent the major cell types found in the small intestine and provide an environment that better mimics the *in vivo* environment. We confirmed the presence of enteroendocrine cells, goblet cells, Paneth cells, enterocytes, and stem cells by gene and protein expression, which aligns with reported data^16,22,32–34^. The presence of tuft cells and M cells was not confirmed here, but other groups have demonstrated them after adding media supplements to promote their differentiation^35,36^. We have observed cell turnover in both formats resembling cell turnover found in the gut *in vivo*, which occurs about every 3 days in the mouse intestine (3-5 in human intestine)^7,27,37–40^. Host cells sloughed off the monolayer and were replaced by new cells in both uninfected and infected conditions. We did not investigate whether turnover rate is affected by infection. Many cells that sloughed off ODMs or enteroids were infected by various stages of *C. parvum* (Sup Fig. 1D). Like many other protozoans, *Cryptosporidium* inhibits host cell apoptosis through NFκB^41–46^. Based on previous studies in biliary epithelium, it may be that uninfected cells in proximity to infected cells are undergoing apoptosis despite prolonged survival of infected cells^47^. A recent study determined that infection did increase cell turnover, but that infected cells were less likely to be apoptotic than neighboring uninfected cells^48^. Mechanisms mediating cell turnover and/or inhibition of apoptosis during *Cryptosporidium* infection are not fully understood, but this system could be helpful in exploring them.

ODMs are ready for infection within 5 days, which more closely resembles the maturation, differentiation, and turnover of cells in the intestine. This is 5-6 days faster than monolayers using the stem cell derived air-liquid interface (ALI) system used in a previously reported *Cryptosporidium* infection model^18^. The defined media used here does not rely on serum supplementation and should not be subject to the same lot-to-lot variation inherent in fetal bovine serum-based media used for the ALI method, and thus be more reproducible. Past studies have demonstrated that serum supplementation may not be sufficient alone to provide the necessary components to support robust oocyst production^14^ or that serum may actually inhibit the infection in some cases^49^.

The main goal of this project was to model successful sexual reproduction leading to production of infectious oocysts in an *in vitro* system, since the programmed nature of *Cryptosporidium’*s lifecycle requires sex for continued infection^10^. Our mating assay confirmed that ODMs support *C. parvum* fertilization *in vitro*. Double positive vacuoles indicated fusion of macrogamonts and microgametes and confirmed that fertilization does indeed occur in ODMs. Double positive vacuoles were likely indicative of zygotes rather than recombined double positive offspring due to the proximity of the mNG and tdT genes on chromosome 5 lowering the probability of recombination resulting in double positive parasites. We would have expected an expansion of the double positive population if successful recombination between those two genes had occurred. This does not mean no recombination occurred, only that it did not occur frequently between the mNG and tdT genes. However, this mating assay method greatly underestimates the number of fertilization events because it fails to reflect the intrastrain fertilization events observed as single positive.

Our fertilization switch reporter is a sensitive tool to measure fertilization that overcomes these shortcomings. Validation of the fertilization switch reporter in mice demonstrated that the color switch is tightly controlled and that rapamycin administration results in a robust shift from tdT to mNG expressing parasites after fertilization occurs. In ODMs, the fertilization switch reporter demonstrated production of mNeonGreen vacuoles of various sizes that were both mononucleated and multinucleated, suggesting they were trophozoites or meronts. Many mCherry positive vacuoles also remained; some appeared to be meronts (multinucleated) and some appeared to be, presumably, unfertilized macrogamonts. If unfertilized macrogamonts can survive and remain in the intestine until another round of sexual differentiation occurs, they could have another chance to be fertilized. The presence of apparent mNeonGreen positive trophozoites and meronts indicates that new oocysts were produced, excysted, and sporozoites infected new host cells. Oocyst excystation has been shown to have a number of triggers, but sometimes only one trigger is required to induce excystation^8^. Changes in temperature, pH, exposure to digestive enzymes and bile acids, and even binding to oocyst wall receptors can trigger excystation^8,9^. It is possible that during media changes oocysts present in culture experienced a temperature change significant enough to trigger excystation. Alternatively, oocyst wall receptors may have bound to host ligands that could have triggered excystation.

Surprisingly, our data using the fertilization switch reporter indicate that the block to fertilization in HCT8 cell culture is not absolute. This result runs counter to a recent mating experiment that detected fertilization *in vivo* but not in HCT8 cell culture using a method in which a Cre recombinase bearing *C. parvum* strain was crossed with a *C. parvum* strain with a floxed termination sequence upstream of a fluorescent reporter gene^12^. In contrast, our data strongly suggest that fertilization and oocyst production, followed by autoinfection occurs in HCT8 cell culture, albeit rarely. The accuracy of this conclusion is supported by the apparent specificity of the fertilization switch reporter *in vivo* and in tissue culture, and by observation of clustered NeonGreen positive vacuoles of various sizes corresponding to different life cycle stages. Interestingly, more mNG clusters were observed after infecting HCT8 monolayers cultured on fibronectin-coated coverslips. As a member of the extracellular matrix, fibronectin could have contributed to enhanced monolayer polarization, but this was not explored^50,51^. The discrepancy with prior work is likely due to enhanced sensitivity of the fertilization switch reporter which theoretically detects all fertilization events compared to previously used methods unable to detect intrastrain mating. Our data are consistent with earlier *Cryptosporidium* research in which production of small numbers of new, usually thin-walled, oocysts in various culture systems was reported, including in HCT8 cell culture^13,14,52^. Nonetheless and consistent with numerous prior reports, the rate of fertilization and oocyst production from HCT8 cell culture was inadequate to support infection of mice, which was only possible with material derived from the new organoid-based culture method. However, our experiments show that HCT8 culture did not support the development of sufficient oocysts to infect mice, which is an inefficient process that would not be detected when coupled with low numbers of oocysts produced in HCT8s. It is also possible that HCT8-derived oocysts are defective. The necessary fertilization enabling factors may be present in HCT8 cell culture but at low levels leading to inefficient and infrequent fertilization. If HCT8s are only capable of supporting production of thin-walled oocysts, those oocysts may not survive passage through stomach acid in the mouse. Explorations into why fertilization and autoinfection occur rarely in HCT8s and more frequently in ODMs and *in vivo* could provide opportunities to develop still simpler continuous culture methods and/or to identify drug or vaccine targets to shorten the duration of infection and prevent transmission.

The programmed nature of the *Cryptosporidium* lifecycle suggests that persistent infection and the disease course are dependent on an appreciable rate of autoinfection. The parasite must need to balance the rate of autoinfection and the rate of release of mature oocysts in the environment in order to find new hosts. In immunocompromised individuals where the infection can become chronic, there may be factors at play that increase the rate of autoinfection. This could be an interesting avenue of research that could benefit the populations most prone to severe cryptosporidiosis.

Proximity of the different strains may be a factor for detecting fertilization via mating experiments using different colored reporters. As has been proposed previously, male and female gamonts develop in close proximity which can facilitate fertilization within the same strain allowing the lifecycle to restart ^10^. Our ability to detect the rare fertilization events in HCT8s using the more sensitive fertilization switch reporter may hinge on this principle.

Whether or to what extent microgametes are motile is still unknown, thus proximity to female gametes may be extremely important for facilitating fertilization. Crossing separate strains may not have detected fertilization due to spacing of initial infection clusters of the different strains and perhaps the sexual stages were not in close enough proximity to facilitate a cross. It may be that *in vivo* more than one complete turn of the life cycle is required before cross fertilization of different strains can occur. Alternatively, the intestinal environment *in vivo* likely provides help in the way of moving microgametes around via peristalsis and flow of luminal contents. Additionally, the tube shape of the intestinal lumen also suggests that parasites may happen upon a new host cell or sexual partner no matter which direction they go, whereas in 2D culture they are limited to a single plane.

Organoid-based methods supporting the complete lifecycle of *Cryptosporidium* have the potential to aid in making big strides toward understanding *Cryptosporidium*’s biology. This system can capitalize on the vast library of transgenic mice to generate organoids with the desired genotype and be used in tandem with genetic manipulation of *C. parvum* to interrogate both sides of the host-parasite interactions^24^. Transfection of the organoids has also been demonstrated by others if the desired host genotype does not exist^53^. Additionally, ODM co-culture with various immune cell types could facilitate *in vitro* studies of immune cell responses to infection^54,55^. Other studies have co-cultured various organoids or ODMs with T cells, intraepithelial lymphocytes, or macrophages^56–60^. Because this method is based on a protocol that makes shifting between host species simple with minor adjustments, it could facilitate the study of the basis of host-parasite specificity^22^. Combining the ODMs with the Fertilization Switch reporter can facilitate studies to identify and verify factors that enable fertilization and to test sexual reproduction inhibitors. Interfering with the parasite’s ability to complete sexual reproduction has the potential to significantly reduce the duration of the infection and reduce associated pathology.

## Materials and Methods

### Media Preparations

L-WRN conditioned media was prepared according to methods previously described^53^. Advanced DMEM/F12 supplemented with 20% FBS was conditioned with stem cell maintenance ligands by culturing L-WRN cells for 12 days. L-WRN cells were engineered to overexpress Wnt3a, R-spondin, and noggin which help maintain and grow intestinal epithelial stem cells. Conditioned media was collected every 4 days and stored at -20°C.

To prepare stem cell maintenance media (SCMM), L-WRN conditioned media was diluted to 50% with Advanced DMEM/F12 and supplemented with 10 mM nicotinamide, 1 mM N-acetylcysteine, 50 ng/mL mouse epidermal growth factor, 1x Pen/strep, 2 mM GlutaMax, 1 mM HEPES, 1X N2 and 1X B27 supplements as described previously^22^. Differentiation media (DM) consisted of Advanced DMEM/F12 supplemented with the same components as SCMM^22^. Five percent SCMM was prepared by diluting 50% SCMM with DM.

### Mouse Intestinal Epithelial Stem Cell Isolation, Culture, and Maintenance

Stem cells were isolated for the small intestinal crypts of an 8 week old female C57Bl6/NJ mouse as described by Miyoshi et al^53^. Briefly, the mouse was euthanized and the small intestine was harvested, flushed with sterile PBS, opened lengthwise, scraped with a sterile glass slide to remove the villi, then minced and submerged in a collagenase solution (2 mg/mL collagenase type 1 and 50 ug/mL gentamicin in DMEM/F12). The tissue fragments were incubated at 37°C with vigorous pipette mixing every 10 minutes for 1 hour until the crypts had separated from larger tissue fragments. Isolated crypts were washed, resuspended in Matrigel™, and dropped on a cell culture plate to form domes. The plate was inverted during polymerization to prevent cell clusters from accumulating at the bottom of the Matrigel™ dome. Once polymerized, SCMM with 10 µM Y27632 inhibitor and 10 µM SB431542 was overlaid. SCMM with 500 nM A83-01 and 1 µM SB202190 added was changed every 2-3 days. For passage, spent media was aspirated, the Matrigel™ domes were washed with PBS, and then overlaid with Trypsin/EDTA and mechanically dissociated with a pipette. Spheroids were incubated with Trypsin/EDTA at 37°C for 2-5 minutes and then washed with washing medium (DMEM/F12, 10% FBS, 2 mM L-glutamine, 15 mM HEPES, and 1x pen/strep). Cells (usually a 1:5 passage) were centrifuged at 200 x g for 5 minutes. The supernatant was removed, and spheroids were resuspended in 2:1 Matrigel™ to SCMM and dotted onto plates to form domes, inverted, and incubated at 37°C until the Matrigel™ polymerized. SCMM was then overlaid on the Matrigel™ domes. Stem cell spheroids were passaged every 3-7 days depending on the size and density of the spheroids.

### Organoid derived monolayer preparation

ODM preparation was adapted from a previously described protocol^22^. Spheroids were expanded appropriately to set up desired experiments. Permeable supports were coated with Matrigel™ diluted 1:10 in SCMM and incubated overnight at 4°C. Prior to seeding, diluted Matrigel™ suspension was removed from wells and permeable supports were allowed to dry at 37°C for 10 minutes. To prepare mISCs for seeding, spheroids were mechanically dissociated in Trypsin-EDTA with vigorous pipetting and incubated for 3-5 minutes at 37°C. Cells were then washed in washing medium, filtered through 40 µm mesh cell strainer, and washed once with PBS. Each permeable support was seeded with 3x10^5^ cells resuspended in 100 µL SCMM supplemented with 10 µM Y27632 inhibitor per well (3x10^5^ cells/permeable support). After 24 hours, media was changed to 5% SCMM with 10 µM Y27632 inhibitor for 24 hours and then switched to differentiation medium without Y27632 inhibitor thereafter. After differentiating for 48 hours ODMs were ready for infection. ODMs were maintained with 5% CO^2^ atmosphere at 37°C. Apical and basal media was changed every 2-3 days. Frequency of media change for apical chambers may have varied due to supernatant collections. New media was added to apical chambers after supernatant collections.

### HCT8 Cell Culture

HCT8s were maintained in RPMI supplemented with 10% FBS, and 1x Pen/Strep. HCT8s were seeded into a 96 well plate or fibronectin coated MatTek plates and infected once confluent. Media was changed every other day. Cells were maintained with 5% CO^2^ atmosphere at 37°C.

### Excystation and Infection

Appropriate numbers of oocysts were disinfected with bleach (10%) for 10 minutes on ice, washed twice with PBS, incubated with 10 mM HCl at 37°C for 10 minutes, centrifuged for 5 minutes at 14000xg, resuspended in 2 mM sodium taurocholate and incubated at 37°C for 10 minutes before infection of ODMs or HCT8s. For transfection, oocysts were allowed to excyst for 45 minutes-1 hour in 2 mM sodium taurocholate at 37°C before transfection. For filtered infections, oocysts were allowed to excyst for 1 hour in 2 mM sodium taurocholate at 37°C before being filtered through a 1 µm syringe filter to remove unexcysted oocysts and retain free sporozoites. Free sporozoites were resuspended in 100 µL/well of DM (ODMs) or RPMI (HCT8s) and added to apical chambers of cell culture supports or to HCT8 cultures. For unfiltered infections, excysted oocysts and sporozoites in sodium taurocholoate were resuspended in warm DM and dispensed into wells in 100 µL volumes. Sporozoites were allowed to infect for 2-3 hours in the incubator at 37°C before washing 3 times with PBS to remove unattached parasites.

### Mouse infection with ODM or HCT8 derived supernatants

HCT8s were maintained in RPMI supplemented with 10% FBS, and Pen/Strep with 5% CO^2^ atmosphere at 37°C. HCT8s were seeded into 24 wells of a 96 well plate and were infected once confluent. This format was chosen to keep the culture surface area comparable between ODM (33.6 mm^2^) and HCT8 culture (33 mm^2^/well). HCT8s and differentiated ODMs were infected with filtered tdTomato *C. parvum* sporozoites (as described above). Supernatants were collected 3 times/week for 2 weeks and stored at 4°C. Media volumes in HCT8s and ODM apical chambers were kept at 100 µL/well throughout the experiment. After 2 weeks, supernatants were spun down at 16000xg for 5 minutes to concentrate any oocysts present and then resuspended in 500 µL of PBS. Concentrated supernatants were gavaged and then used to gavage 5 mice/condition (IFNg-/-, 3-4 weeks old Male). Infection was monitored via the nanoluciferase assay (Promega).

## Acknowledgements

Funding for this study was provided by the Vermont Immunobiology and Infectious Disease Training Grant 2T32AI055402-16A1 and RO1AI143951 to CDH.

We would like to thank the Harry Hood Bassett Flow Cytometry and Cell Sorting Facility (RRID:SCR_022147) at the University of Vermont Larner College of Medicine in the generation of our flow cytometry data.

Transmission electron microscopy sample preparation and imaging was performed at the Microscopy Imaging Center at the University of Vermont (RRID# SCR_018821).

The qPCR and qRT-PCR were performed using equipment in Vermont Integrative Genomics Resource DNA Facility and was supported by University of Vermont Cancer Center, Lake Champlain Cancer Research Organization, and the UVM Larner College of Medicine.

We would like to thank Dr. Christian Klotz for consultations while refining this protocol for *Cryptosporidium* infection, Dr. Boris Striepen for the generous gift of the tdTomato and mNeonGreen strains of *C. parvum*, and Camilla Strother and Ethan Mattice for their help during the initial stem cell isolations.

Figure 1A, 2A, and 3A were created with BioRender.com.

