## Supplemental Methods and Figures for "Organoid-based *in vitro* system and reporter for the study of *Cryptosporidium parvum* sexual reproduction"

**Microscopy:** Imaging was performed using a Nikon widefield epifluorescence microscope equipped with a 1.4-megapixel EXi blue CCD camera. For live imaging studies, infected cultures were briefly removed from the incubator for periodic image collection at specified timepoints throughout the infection. A random sampling of tiled images (3x3-5x5) to cover approximately 10% of the permeable support's surface area were collected at 20x. Images were analyzed visually or parasitic vacuoles were counted using a macro for Image J. Vacuoles counted for imaged area were then used to estimate the total number of vacuoles present on the total cell culture surface area. Deconvolution was done using AutoQuant software. Imaging was performed on a Nikon widefield epifluorescence microscope equipped with a 1.4-megapixel EXi blue CCD camera.

**Immunofluorescence assay**: Samples were fixed with 4% formaldehyde for 15 minutes at room temperature and then washed with PBS. Samples were permeabilized with 0.25% Triton X 100 in PBS for 10 minutes and then washed with PBS and blocked with 4% BSA in PBS overnight at 4°C. For experiments with Fertilization Switch *C. parvum*, ODMs were washed with TBS then incubated with anti-mNeonGreen at 1:200 for 1 hour at RT or overnight at 4°C. ODMs were washed 3 times with PBS with 0.1% Tween 20 and then incubated with goat anti-mouse AF488 1:300 for 1 hour at RT. ODMs were washed and blocked for 1 hour at RT, incubated with anti-RFP antibody overnight at 4°C, then washed and stained with goat anti-Rabbit AF568 at 1:300 for 1 hour at RT, then washed 3x with with PBS with 0.1% Tween 20 and counterstained with 20 µg/mL Hoechst for 10 minutes at RT. For staining for epithelial cell type markers, samples were incubated with primary antibody at 5-10 ug/mL overnight at 4°C, washed with 4% BSA, then secondary antibody was added at 2 ug/mL and incubated at room temperature for 1 hour, then samples were washed and counterstained with 20 µg/mL Hoechst for 10 minutes. Stained ODMs were cut out of the supports with a razor blade and mounted on glass slides with Fluoro-gel mounting medium.

**Nanoluciferase assay:** The assay was performed according to a previously described protocol[61]. A single fecal pellet per mouse per day was homogenized using glass beads in lysis buffer. Samples were allowed to lyse for 20 minutes before adding nLuc substrate to the sample. Samples were plated in white opaque 384 well plates skipping every other well and were read on a BioTek plate reader.

**Generation of genetically modified *C. parvum*:**

**Plasmid Construction:** All G blocks except for the block containing the HAP2 promoter were synthesized by Twist Biosciences (see supplemental materials list). To generate a G block for the HAP2 promoter, the region 339 bp upstream of HAP2 gene was PCR amplified from *C. parvum* IOWA genomic DNA (see supplemental materials list). Gibson assembly failed with the original G-blocks for the Fertilization Switch construct. To reduce the number of G-blocks used for the Fertilization Switch construct, new G-blocks were designed and ordered to reduce the number of G-blocks. However, G-blocks could not be designed for one region to combine previously designed G-blocks. To create an in-house G-block 8, G-blocks 7, 8.1, 8.2, 8.3, and 9 were combined using Gibson assembly. The resulting fragment was used as template and a portion of it was PCR amplified to create a single G-block. Plasmids with a Puc19 backbone were constructed using the GeneArt™ Gibson assembly HiFi Cloning Kit and appropriate G-blocks for the fertilization switch constructs. *E. coli* was transformed with the Fertilization Switch plasmid, clones were screened by restriction digest, and candidate clones were verified by sequencing via Plasmidsaurus. Selected clone plasmids and the CRISPR/Cas9 plasmid were grown up in *E. coli* and DNA was purified using a DNA midi prep (Promega). DNA was concentrated by ethanol precipitation. Both constructs were inserted into the thymidine kinase locus (cgd5_4440).

**Generation of Transgenic parasites: CRISPR/Cas Transfection:** Excysted *C. parvum* sporozoites were transfected with 50 μg Cas9 plasmid and 50 μg linearized repair cassette in SF Buffer (Lonza) using the EH100 program on the Lonza 4D Nucleofector Nucleofection System according to a previously described protocol [23,61]. Transfected sporozoites were resuspended in sterile PBS and blue food coloring and kept on ice until injected into the small intestine of IFNγ-/- mice.

**Surgical infection of mice with transgenic parasites**. Four IFNγ-/- mice (3-4 weeks) were surgically infected with transfected *C. parvum* sporozoites according to a previously described protocol [23]. Briefly, mice were anesthetized with isofluorane in a chamber then moved to a sterile surgical area with isofluorane delivered through a nose cone. A medial cut was made through the skin and peritoneum just under the sternum in the abdomen. A small loop of the small intestine was accessed and transfected sporozoites were injected into the small intestinal lumen. The intestine was returned to the abdominal cavity and the peritoneum and skin were sutured closed. Animals were monitored for recovery from anesthesia and healing at the surgical site. Fecal samples were collected and monitored for successful infection using the Nano-Glo luciferase assay (Promega) according to a previously described protocol [23,61]. The mice were maintained under paromomycin selection (20 g/L) in drinking water starting the day after surgery. Fecal oocysts were purified by sucrose gradient followed by cesium chloride gradient. Some oocysts were used to infect NSG mice for maintenance of the transgenic lines. Fecal collections were performed using fecal collection cages with damp Kimwipes™ lining the cage bottom underneath metal grates keeping the mice elevated above the cage floor to preserve the fecal pellets. Infection was monitored by nLuc assay.

**Animal use:** mNeonGreen and tdTomato *C. parvum* oocysts were maintained by infecting Nod/SCID/Gamma mice aged 3-6 weeks old with $10^5$ mNeonGreen or tdTomato oocysts that were a kind gift from the lab of Dr. Boris Striepen. Paromomycin was added to drinking water (16 mg/mL) and offered to mice ad libitum to select for mNeonGreen and tdTomato parasites. Fecal samples were collected for up to 3 months. Oocysts were isolated and purified from fecal samples using a sucrose floatation and cesium chloride gradient[52]. A C57Bl6/NJ (female, 8 weeks) mouse was euthanized and  mouse intestinal stem cells were isolated from the ileum. Four IFNg-/- mice (3-4 weeks) were surgically infected with transgenic FS *C. parvum*. Animal experimentation and use was approved by the Institutional Animal Care and Use Committee of the University of Vermont (protocol X0-153).

***In vivo* Fertilization Switch reporter validation**: NSG mice infected with Fertilization Switch reporter *C. parvum* were separated into 2 cages. One cage contained one mouse that was left untreated. The other cage contained 2 mice that both received rapamycin 3 times/per day in 200 µL doses for a total of 10 mg/kg/day. Fecal collections were performed as described above 1 day before and 7 days after the rapamycin treatment period.

**Oocyst midiprep**: About 500 mg of fecal sample was placed in a 15 mL conical tube and homogenized by vortexing in 2-5 mL of tap water. After allowing the large particulates to settle for 2-5 minutes the supernatant was transferred to a fresh conical tube. This was repeated until the supernatant was almost clear after vortex mixing and particulate settling. Collected supernatant spun down for 5 minutes at 3000xg and resuspended in 5 mL tap water and overlaid onto 5 mL of sucrose solution (1.33 specific gravity), mixed by inversion and centrifuged at 1000xg for 5 minutes. Supernatant was collected and divided in 2 mL microcentrifuge tubes, 1 mL each. One mL of tap water was added to each tube, mixed, and spun down at 10,000xg for 5 minutes. Supernatant was discarded and pellets were combined by resuspending in 1 mL tap water, then overlaid on 1 mL 1.25 M cesium chloride and spun down for 3 minutes at 16,000xg. The entire supernatant was collected, distributed into 2 fresh tubes, diluted with 1 mL water, and spun down for 3 minutes at 16,000xg. Pellet was washed twice and oocysts were resuspended in 1 mL PBS and stored at 4°C. This procedure was adapted from a procedure described by Sebastain Shaw. Purified fecal oocysts were used for microscopic analysis, qPCR, and flow cytometry.

**RNA extraction and qRT-PCR:** RNA extraction and purification from tissue culture was performed using RNeasy following manufacturer's instructions. RNA concentration was quantified, normalized, and then converted to cDNA using ThermoScientific High-Capacity cDNA kit. Expression of genes of interest (ChgA, Muc2, ZO-1) was assessed using SYBR Green PowerUp Mastermix and mRNA relative expression was normalized to GAPDH and calculated using the ΔΔCT method.

**qPCR/PCR**: DNA collection and purification from organoid derived monolayers (ODMs) and oocysts using the QIAmp DNA mini Kit. ODMs were scratched and mixed by pipetting in ATL Buffer and proteinase K for lysis and enteroids were collected by centrifugation and resuspended in ATL Buffer and proteinase K. Samples were incubated at 56°C for 16-24 hours and then proceeded to column purification using manufacturer instructions. A standard curve was generated from a dilution series of purified *C. parvum* (Bunchgrass) DNA. Primer sequences for *C. parvum* 18S rRNA (forward: 5'-TAGAGATTGGAGGTTGTTCCT-3'; reverse: 5'-

CTCCACCAACTAAGAACGGCC-3') were used for quantification. qPCR reactions were performed with *Power*SYBR Green, 2 µL of DNA template on a QuantStudio6 using these amplification conditions: 95°C 10 min, 40 cycles of 95°C 15 sec, 60°C 60 sec and a melt curve analysis. Genomic equivalents were determined by comparison to the standard curve. mCherry excision was verified by PCR amplification using upstream and downstream primers (Table 3.1) outside of the mCherry region of the construct. PCR products were run on a 1% agarose gel with ethidium bromide and read with a GelDoc (BioRad). DNA Ladder used was 1 kb GeneRuler (ThermoFisher Scientific).

**Flow cytometry**: Supernatants collected from ODMs or purified fecal oocysts were fixed in 4% formaldehyde for 15 minutes, permeabilized with Triton X 100 for 10 minutes, blocked with 4% BSA for 1 hour at RT, and stained with 5 µg/mL anti-COWP antibody overnight at 4°C. Samples were washed and stained with 2 µg/mL rabbit anti-mouse AF633 secondary antibody for 1 hour at RT. Samples were washed and resuspended in 1 mM EDTA in PBS and used for flow cytometry analysis on the MACSQuant VYB or Cytek Aurora. Flow cytometric data analysis was done using FlowJo Software. Oocysts in supernatants were quantified using FSC/SSC gates to identify single cell populations of oocyst size by comparing to stock oocysts and identifying COWP+ cells. Then COWP+ cells were assessed for mNeonGreen and/or mCherry expression.

**Transmission Electron Microscopy**: For preparation of ODMs, media from the apical and basal chambers was removed and replaced with Karnovsky's fixative (2.5% glutaraldehyde, 1% paraformaldehyde in 0.1M cacodylate buffer). Samples were fixed for 2 hours or overnight at 4°C. ODMs were washed with 0.1M cacodylate buffer. Samples were then post-fixed in 1% $OsO_4$ in 0.1M cacodylate buffer for 45 min at 4ºC, rinsed with cacodylate buffer, dehydrated in a graded ethanol series and embedded in Spurr's resin. Ultrathin sections were cut with a diamond knife, retrieved onto copper grids, and contrasted with uranyl acetate and lead citrate. Finally, the sections were imaged on a JEOL 1400 transmission electron microscope, and digital images acquired with an AMT XR611 high resolution 11-megapixel mid-mount CCD camera.

**Statistical analysis:** GraphPad Prism was used for all statistical analyses. Statistical significance was assessed with a two-tailed, unpaired t-test or One-Way ANOVA where p value $\leq 0.05$ was considered significant. Error is shown as standard deviation and p-values are displayed on each graph. Statistical parameters are specified in figure legends.

**Data Availability:** Data are in the form of images and excel files with counts generated by macro analysis in Image J, qRT-PCR, or flow cytometry data. All data will be made available upon request via file transfer.

### Supplemental Materials List

| Material | Vendor | Product # |
|---|---|---|
| Advanced DMEM/F12 | Gibco | 12634028 |
| Nicotinamide | Millipore | 481907 |
| N-acetylcysteine | Sigma | A9165 |
| mouse epidermal growth factor | R&D Systems | 2028-EG |
| Penicillin/streptomycin | Corning | 30-002-Cl |
| GlutaMax | Gibco | 35050061 |
| HEPES | Gibco | 15630-080 |
| N2 supplement | Gibco | 17502048 |
| B27 supplement | Gibco | 17504044 |
| Matrigel™ | Corning | CB40234 |
| Y27632 inhibitor | R&D Systems | 125450 |
| SB 431542 | R&D Systems | 16141 |
| A83-01 | Sigma | SML0788 |
| SB 202190 | SYNthesis | SYN-1073 |
| 0.25% Trypsin/EDTA | Gibco | 25200-056 |
| High-Capacity RNA-to-cDNA Kit | Applied Biosystems | 4387406 |
| RNeasy | QIAGEN | 74104 |
| PowerUp Mastermix | Applied Biosystems | 4367659 |
| QIAmp DNA mini Kit | Qiagen | 51306 |
| ThinCert Tissue Culture inserts | Greiner Bio-one | 662 641 |
| Fetal bovine serum, H.I. | Sigma | 12306C, Lot 17H093 |
| Fluorogel | Electron Microscopy Science | 17985-10 |
| Prolong Antifade Gold | Invitrogen | 936941 |
| EDTA (pH 8.0) | Promega | V4231 |
| 4% Paraformaldehyde | ThermoScientific | J19943-K2 |
| DAPI | AppliChem | A4099, 0010 |
| Hoescht | AnaSpec Inc. | AS-83219 |
| Bovine Serum Albumin | Sigma | A7906 |
| Triton-X-100 | Promega | H5142 |
| Tween 20 | Fisher Bioreagents | BP337-500 |
| MicroAmp Optical Reaction plates | Applied Biosystems | 4309849 |
| 1.2 µm Syringe filter | Sartorius | 17593 |
| 40 µm nylon Cell strainer | BD Falcon | 352340 |
| GeneArt™ Gibson assembly HiFi Cloning Kit | ThermoFisher Scientific | A46626 |

| Cell lines | | |
|---|---|---|
| L-WRN | ATCC (Miyoshi et. al 2013) | CRL-3276 |
| C57Bl6/NJ intestinal stem cells | Isolated in Huston lab | NA |
| HCT8 | | |

| Equipment | | |
|---|---|---|
| MACSQuant VYB | Miltenyi Biotec | Harry Hood Bassett Flow Cytometry and Cell Sorting Facility, UVM |
| CytekAurora | Cytek | Harry Hood Bassett Flow Cytometry and Cell Sorting Facility, UVM |
| QuantStudio6 | Applied Biosystems | Vermont Integrative Genomics Resource DNA Facility, UVM |
| JEOL 1400 transmission electron microscope | JOEL | Microscopy Imaging Center, UVM |
| Lonza 4D Nucleofector Nucleofection System | Lonza | NA |
| | Nikon | NA |

| Parasites | |
|---|---|
| *C. parvum* | Bunchgrass Farms |
| tdTomato *C. parvum* | gift from Striepen Lab |
| mNeonGreen *C. parvum* | gift from Striepen Lab |

| Software | |
|---|---|
| ImageJ 1.53k | NIH |
| Adobe Illustrator | Adobe |
| FlowJo 10.8.0 | BD |
| QuantStudio 1.3 | Applied Biosystems |
| MS Excel 2010 | Microsoft |
| GraphPad Prism 10 | GraphPad |

| Antibodies/stains | Host | Source | Product # |
|---|---|---|---|
| Anti-ChgA | mouse | Proteintech | 60135-1-Ig |
| Anti-Lysozyme | rabbit | Proteintech | 15013-1-AP |
| Anti-Muc2 | rabbit | Proteintech | 27675-1-AP |
| Anti-ZO-1 | mouse | Invitrogen | 33-9100 |
| Anti-COWP1 | mouse | Invitrogen | MA183095 |
| anti-Mouse, Alexa Fluor 488 | Goat | LifeTechnologies | A11029 |
| anti-Rabbit, Alexa Fluor 488 | Donkey | LifeTechnologies | A21206 |
| Alexa Fluor 633 phalloidin | NA | Invitrogen | A22284 |
| FITC-VVL | NA | Vector Labs | FL-1231 |
| Anti-mNeonGreen | Mouse | Chromotek/Proteintech | 32F6 |
| Anti-RFP | Rabbit | Rockland Immunochemicals | 600-401-379 |

| | Target | Sequence 5'-3' | PrimerBank ID/Reference |
|---|---|---|---|
| housekeeping | mGapdh FWD | AGGTCGGTGTGAACGGATTTG | 6679937a1 |
| | mGapdh REV | TGTAGACCATGTAGTTGAGGTCA | |
| stem cell | mLgr5 FWD | CCTACTCGAAGACTTACCCAGT | 6753842a1 |
| | mLgr5 REV | GCATTGGGGTGAATGATAGCA | |
| enteroendocrine | mChgA FWD | ATCCTCTCTATCCTGCGACAC | 6680932a1 |
| | mChgA REV | GGGCTCTGGTTCTCAAACACT | |
| goblet cell | mMuc2 FWD | GCTGACGAGTGGTTGGTGAATG | Holthaus et al., 2021 |
| | mMuc2 REV | GATGAGGTGGCAGACAGGAGAC | |
| tight junction | mZo1 FWD | GCCGCTAAGAGCACAGCAA | 6678355a1 |
| | mZo1 REV | TCCCCACTCTGAAAATGAGGA | |
| *C. parvum* | 18S rRNA FWD | TAGAGATTGGAGGTTGTTCCT | NA |
| | 18S rRNA REV | CTCCACCAACTAAGAACGGCC | |

**Primers**

| Purpose | Sequence 5'-3' | PrimerBank ID/Reference |
|---|---|---|
| For mCherry excision assessment | Cre60 FWD | CTATTTATTCTGCCGGGTCAG |
| | cgd1_3020 REV | GAGATTTTTCACCAATTTCTACTTTATC |
| For G block 8 creation | Cre60 FWD | CTATTTATTCTGCCGGGTCAG |
| | nluc REV | ATCACCTTAAAGTGATGATC |

**FS G-blocks**

| | G-blocks for Gibson Assembly | | |
|---|---|---|---|
| | **Name** | **Length (bp)** | **Sequence** |
| **Fertilization Switch** | **GB_12** | 1075 | CAGGAAACAGCTATGACCATGATTACGCCAAGCTTGCATGCCTGCAGGTCT<br>TTAAAGCAATAATATCACTCATACCTACTGCAAATAAGATTGGAAATACGC<br>TAGGAATTAAGATAAAAAGAAAAACTTAATCGATACTATCCTACACGCCAC<br>GAACTTATTATTAATTCAATTTACACCACGCAGCCCAGGATCTGCATACAGA<br>TAATAACATTTTCCATGTATGTTCAGAAAATGCTTGCCTGTCCCTAAACATG<br>TCCATCAGATTTTTCCTGACTTCATCTGAGGTGGATCCATTGCTAGCACCCG<br>GGTTGCTAGGTGAAGCTTCTAATTTTAGAAGCTCTACATCAAAGACGAGAG<br>TTGCATGTGGTGGAATTATACCTGGGTGTCCAGTGGCACCATAAGCATAAT<br>CTGGAGATATAGTCAATTTTGCTCTTTGTCCTACACTCATTTGAGCAACTCC<br>TTCTTCCCAGCCTCGAATAACCTCTTGCTTGCCTAGCATAAATTTAAAGGGC<br>TTATTTCTGTCCCGACTTGAATCGAATTTTTTTCCATCTTCAAGCATCCCGGT<br>GTAATGCACAACACATGTCTGGCCACGTTTAGGAAACGTGCGCCCATCTCC<br>TGGCGAGATTGTTTCAACTTGTACTCCTCTAGAAACTTTTCTTTTCTTTTTAG<br>GTGCCATGGTGGCAATAAAAGAAAAAACTATTTAATGAAATTGACCTAATA<br>ATTAAATTAGGGCAGATATTCAAATAGTAATTGAAATTATTAGGTAAAAAA<br>TTGTTGATTGAGGAAAAACAATAATTCGAGGATACACCTAACGCATATACG<br>ATACAATTTACACAAATTCGTGAGTGCTCATATTAGAGTTTAATCTAAATTG<br>TTTTAAAGAAACATATGATAATAAACTATCGCTTTGCAATAAATTAAAATAT<br>TGAACTTTCGTTGGAATGTGTTTTTTCAATAATTTTTGATCGGATTGTAAATT<br>TATTGGATAAAACCAATCAATTTTTTGGGCGCAATTTTCAGGAAAATCAAA<br>CTATTAATTTAGAGATTGATTAAATAATTTTTAATTGACATA |
| | **GB_13** | 1186 | ATTAAGAGGCACTAATTTTCCTTCCGATGGGCCCGTAATGCAAAAAAAGAC<br>AATGGGATGGGAAGCATCATCCGAGCGGATGTATCCAGAGGATGGGGCAC<br>TGAAAGGCGAAATTAAACAAAGGTTAAAACTTAAGGATGGAGGACATTAC<br>GATGCTGAAGTCAAAACCACATACAAAGCAAAAAAGCCTGTTCAGCTACCA<br>GGTGCATATAACGTCAATATAAAATTGGATATCACCAGCCACAATGAAGAT<br>TATACAATAGTAGAACAATATGAAAGAGCAGAGGGTAGACATTCAACTGGT<br>GGGATGGATGAATTGTATAAATAAATAACTTCGTATAGCATACATTATACG<br>AAGTTATGCTAGCATGGTTAGTAAAGGAGAAGAAGACAATATGGCAAGCTT<br>GCCAGCTACCCATGAACTTCACATTTTTGGATCAATTAATGGTGTTGATTTT<br>GATATGGTTGGTCAAGGAACTGGAAACCCAAACGACGGTTACGAAGAACTT<br>AATTTAAAATCAACTAAAGGTGATTTGCAATTCTCACCTTGGATATTAGTAC<br>CACATATTGGGTACGGATTCCATCAATATTTACCTTATCCAGATGGGATGTC<br>TCCATTCCAAGCAGCAATGGTCGACGGATCAGGATATCAGGTACATAGAAC<br>CATGCAATTTGAGGACGGTGCATCACTAACAGTTAATTATAGATATACTTAT<br>GAAGGCAGTCACATTAAGGGCGAAGCACAAGTTAAAGGGACAGGTTTCCC<br>AGCCGATGGTCCAGTTATGACTAATTCTTTAACTGCCGCTGATTGGTGTCGA<br>TCTAAAAAAACATATCCTAATGATAAGACTATTATTTCAACTTTTAAATGGT<br>CATATACTACAGGCAACGGAAAAAGATATAGGAGTACAGCTAGGACTACTT<br>ATACTTTTGCAAAACCAATGGCTGCCAATTACTTAAAAAATCAACCGATGT<br>ATGTTTTCCGTAAAACGGAATTGAAACACTCAAAAACTGAATTGAATTTCA<br>AAGAATGGCAGAAAGCATTTACTGACGTTATGGGAATGGATGAATTATATA<br>AATAATAATAAGTTCGTGGCGTGTAGGATAGTATCGATTAAGTTTTTCTTTT<br>TATCTTAATTTGGGGAAACTAAATATACTGAAATTCGGTAGATTCTATATCT<br>CACGGGAC |
| | **GB_3** | 598 | TAATAAGTTCGTGGCGTGTAGGATAGTATCGATTAAGTTTTTCTTTTTATCTT<br>AATTTGGGGAAACTAAATATACTGAAATTCGGTAGATTCTATATCTCACGG<br>GACAGCTTTTCACACACACTTAGTTCTATATTGCGTCATAACTTTTGTTTTTT<br>TTTGTTGCACTTTTTTTCTCTTATATTCAAGTAAGTGGTTTAGATTCTCTAAG<br>GGCGGGAATATGAATTAGTGGCAATTCAAAGGATTTAAACGATCAGGCGCC<br>TGCACACCAAACTCTAATTCGTACACAGCATGCCCGAGGTTATAGATATAG<br>ATACTGCGAAATTATTTTCATTGTTTCAGTTAAGAAATAATAAAATATTTTA<br>TTATAGTTATTTTCCAAATTTATTTGAGATTTTTGTATTGAAGTTTAGCCGTC<br>GACATGGTCTTCACACTCGAAGATTTCGTTGGGGACTGGCGACAGACAGCC<br>GGCTACAACCTGGACCAAGTCCTTGAACAGGGAGGTGTGTCCAGTTTGTTT |

| | | | |
|---|---|---|---|
| | | | CAGAATCTCGGGGTGTCCGTAACTCCGATCCAAAGGATTGTCCTGAGCGGT<br>GAAAATGGGCTGAAGATCGACATCCATG |
| | **GB_5** | 1355 | CCAAAGGATTGTCCTGAGCGGTGAAAATGGGCTGAAGATCGACATCCATGT<br>CATCATCCCGTATGAAGGTCTGAGCGGCGACCAAATGGGCCAGATCGAAAA<br>AATTTTTAAGGTGGTGTACCCTGTGGATGATCATCACTTTAAGGTGATCCTG<br>CACTATGGCACACTGGTAATCGACGGGGTTACGCCGAACATGATCGACTAT<br>TTCGGACGGCCGTATGAAGGCATCGCCGTGTTCGACGGCAAAAAGATCACT<br>GTAACAGGGACCCTGTGGAACGGCAACAAAATTATCGACGAGCGCCTGATC<br>AACCCCGACGGCTCCCTGCTGTTCCGAGTAACCATCAACGGAGTGACCGGC<br>TGGCGGCTGTGCGAACGCATTCTGGCGGCTAGCATGATTGAACAAGATGGT<br>TTACACGCTGGTTCTCCCGCCGCTTGGGTCGAAAGACTTTTCGGTTATGACT<br>GGGCTCAACAAACCATCGGTTGCTCTGATGCCGCCGTCTTCCGTCTTTCTGC<br>TCAAGGTCGTCCTGTTCTTTTCGTCAAGACCGACCTTTCTGGTGCCCTTAAT<br>GAACTTCAAGATGAAGCTGCCCGTCTTTCTTGGCTTGCCACCACCGGTGTTC<br>CTTGCGCTGCTGTCCTTGACGTTGTCACTGAAGCCGGTAGAGACTGGCTTCT<br>TTTAGGTGAAGTCCCCGGTCAAGATCTTCTTTCTTCTCACCTTGCTCCTGCCG<br>AAAAAGTTTCTATCATGGCTGATGCTATGCGTCGTCTTCATACCCTTGATCC<br>CGCTACCTGCCCTTTCGACCACCAAGCCAAACATCGTATCGAACGTGCTCGT<br>ACTCGTATGGAAGCCGGTCTTGTCGATCAAGATGATCTTGACGAAGAACAT<br>CAAGGTCTTGCCCCTGCCGAACTTTTCGCCAGACTTAAGGCCCGTATGCCCG<br>ACGGTGAAGATCTTGTCGTCACCCATGGTGATGCCTGCTTACCCAATATCAT<br>GGTTGAAAATGGTCGTTTTTCTGGTTTCATCGACTGTGGTCGTCTTGGTGTC<br>GCCGACCGTTATCAAGATATTGCCTTAGCTACCCGTGATATTGCTGAAGAAC<br>TTGGTGGTGAATGGGCTGACCGTTTCCTTGTCCTTTACGGTATCGCCGCTCC<br>CGATTCTCAACGTATCGCCTTCTATCGTCTTCTTGACGAATTCTTCTGATAAT<br>AAGTTCGTGGCGTGTAGGATAGTATCGATTAAGTTTTTCTTTTTATCTTAATT<br>TTAATTAAGGCAAAATTTGGCGCAGCTATGGCGCCTCTTGAAAGTGCCTAA<br>AAAGGAGGACTCTAGAGGATCCCCGGGTACCGAGCTCGAATTCACTGGCCG<br>TCGTTTTACA |

| | G-blocks for Gibson Assembly | | |
|---|---|---|---|
| | **Name** | **Length (bp)** | **Sequence** |
| **Fertilization Switch** | **GB_7** | 1256 | TGAGAGAAAATTTAAAAATTTAAGATGAAAGAAGAATAAAATAATATATAGCC<br>ACCATGGCACCAAAAAAGAAAAGAAAAGTTTCTAGAATACTGTGGCATGAAAT<br>GTGGCATGAAGGCTTGGAAGAGGCATCTCGTTTGTATTTTGGGGAAAGGAATGT<br>AAAAGGAATGTTTGAGGTTTTAGAACCGTTGCATGCTATGATGGAACGGGGAC<br>CCCAAACTTTAAAAGAAACATCATTTAATCAGGCATATGGTCGAGATTTAATGG<br>AAGCACAAGAGTGGTGTAGGAAATATATGAAATCAGGAAATGTCAAGGATCTA<br>CTACAAGCGTGGGATCTATATTATCATGTATTCCGACGAATATCAGCTAGCCCC<br>AGCAACCCTGGCGCTAGCAATGGATCCAATAGGAAATGGTTCCCTGCTGAACC<br>AGAGGATGTAAGGGATTACCTATTGTATTTACAAGCAAGAGGACTTGCTGTTAA<br>AACGATACAACAGCACTTGGGCCAGCTAAACATGTTGCATAGGAGAAGTGGAT<br>TACCAAGACCTTCTGATTCAAATGCTGTTTCCCTTGTGATGAGGAGAATAAGAA<br>AAGAAAATGTTGATGCTGGAGAGAGAGCAAAACAAGCTTTGGCATTTGAACGC<br>ACTGATTTTGACCAAGTCAGATCATTAATGGAGAATTCTGATAGATGTCAGGAT<br>ATCAGGAACCTCGCATTCTTGGGAATTGCCTACAATACTTTGTTAAGAATTGCA<br>GAAATTGCAAGAATTAGAGTGAAAGATATATCCCGCACAGATGGAGGAAGAAT<br>GTTAATCCATATTGGCAGGACTAAGACACTTGTTTCAACAGCTGGTGTGGAAAA<br>AGCATTATCCCTTGGGGTTACTAAATTAGTTGAAAGATGGATAAGTGTTTCTGG<br>AGTAGCTGATGACCCGAATAACTATTTATTCTGCCGGGTCAGAAAAAATGGTGT<br>AGCTGCACCAAGTGCCACCTCACAATTATCCACCCGGGCATTAGAAGGAATATT<br>TGAGGCAACACACCGCCTTATTTATGGAGCAAAAGATGACTCTGGACAAAGAT<br>ATTTAGCATGGTCTGGACATAGTGCAAGAGTAGGTGCTGCCAGGGACATGGCC<br>AGGGCTGGTGTGTCGATCCCAGAAATTATGCAAGCTGGTGGCTGGACTAATGTT<br>AATATTGTAATGAACTACATTAGAAATTTGGACTCTGAGACTGGGGCCATGGTT<br>AGGTTGCTAGAGGATGGGGACTAA |
| | ***GB_8*** | 1043 | CTATTTATTCTGCCGGGTCAGAAAAAATGGTGTAGCTGCACCAAGTGCCACCTC<br>ACAATTATCCACCCGGGCATTAGAAGGAATATTTGAGGCAACACACCGCCTTAT<br>TTATGGAGCAAAAGATGACTCTGGACAAAGATATTTAGCATGGTCTGGACATA<br>GTGCAAGAGTAGGTGCTGCCAGGGACATGGCCAGGGCTGGTGTGTCGATCCCA<br>GAAATTATGCAAGCTGGTGGCTGGACTAATGTTAATATTGTAATGAACTACATT<br>AGAAATTTGGACTCTGAGACTGGGGCCATGGTTAGGTTGCTAGAGGATGGGGA<br>CTAATAATAAGTTCGTGGCGTGTAGGATAGTATCGATTAAGTTTTTCTTTTTATC<br>TTAATTAATACTAATTCTTTTGAATTTCCACATATCTAATCATAATACTAGTCTC<br>TAACATATAAGACTCATTTCTCTGGAGAAAGGTACATATATATTGGAGCAATTC<br>ATTATGGGATTCACACTCCCTCTACCCAATATTATCAATAATACAGATAATCAA<br>CAATTAGTAGTAGCATTTCTAAATGGTCTACTAGCAGGAATATGTCATTAAATT<br>TTCTAACAATCTATTAAATTATTGTGATTCTCTAAGATATTTGTGTTCCACTTAT<br>TTTATTAAAAACTAATATTTTTACTTGGAAGTATTTTCTTTTGCTATTTCTTAACT<br>CAGATATACCATTCCCTACGACTTGTGAAATATTTTACTAATACTTAACTAAAA<br>CTATACAAGATTTTTTCCCACTCAAGTTCTAATCAATGCTTACTTTAAACACGTT<br>TTTTTACACCTATATATACTTTAATATTTGTTTTCAATATTTTTATTTATTTGCAT<br>GCAAATTATATATCTAAAAATGGCGCCACAATGGCTGGTATATAATATCCAGCC<br>TGCGAGCATGCTTCAGATTTAAAGGAAAAAGGGAGAAACAGCACAAAATTTGT<br>ACTTGTTAGAATCGCAAATAAATTTTGTTAAAAAAATTTGATAAAGTAGAAATT<br>GGTGAAAAATCTC |
| | **GB_9** | 517 | AAAGTAGAAATTGGTGAAAAATCTCATTTTATTTGTTTCAAAAGAAAAAAATAA<br>CTTCGTATAGCATACATTATACGAAGTTATATGGTTAGCAAAGGAGAAGAGGA<br>TAACATGGCAATTATAAAAGAATTCATGAGATTTAAGGTTCATATGGAAGGTTC<br>TGTAAATGGACACGAGTTTGAAATTGAAGGAGAAGGTGAAGGAAGGCCATACG<br>AAGGGACACAAACTGCAAAATTAAAAGTTACTAAAGGTGGACCTCTCCCATTC<br>GCATGGGATATTTTATCTCCTCAATTTATGTATGGATCTAAGGCATACGTCAAA<br>CATCCAGCAGATATCCCGGATTACTTGAAACTTAGTTTTCCAGAAGGTTTTAAA<br>TGGGAAAGGGTTATGAATTTCGAGGACGGAGGTGTGGTTACTGTGACGCAAGA<br>TTCAAGTCTACAGGATGGTGAATTTATTTATAAGGTAAAATTAAGAGGCACTAA<br>TTTTCCTTCCGATGGGCCCGTAATGCAAAAAAAGA |

| | | | |
|---|---|---|---|
| | **HAP2 promoter** | 339 | GAAAATCAAACTATTAATTTAGAGATTGATTAAATAATTTTTAATTGACATAAA<br>GTTTGTTTAAAGAATATTATTAAAAAATAAGGAATTGGATAAAGTGATATGATG<br>AGTATTTAACAGTATCAGCTAAAATAATTAAAGTTATGGAGGAAAATCTATAA<br>ATATTTTTATTTGTTAATATGACCACACAAATAATAAGAGAATAATTAAAAAGT<br>AAAATAGTGTGGGAGTTAAAAAAAGAATAAGTCGTTATGAAAGATATATAAAT<br>TTATATGTAATGGTTTAAAATTGAGAGAAAATTTAAAAATTTAAGATGAAAGA<br>AGAATAAAATAATATATA |
| | ***GB_8.1*** | 387 | GGCTGGACTAATGTTAATATTGTAATGAACTACATTAGAAATTTGGACTCTGAG<br>ACTGGGGCCATGGTTAGGTTGCTAGAGGATGGGGACTAATAATAAGTTCGTGG<br>CGTGTAGGATAGTATCGATTAAGTTTTTCTTTTTATCTTAATTAATACTAATTCT<br>TTTGAATTTCCACATATCTAATCATAATACTAGTCTCTAACATATAAGACTCATT<br>TCTCTGGAGAAAGGTACATATATATTGGAGCAATTCATTATGGGATTCACACTC<br>CCTCTACCCAATATTATCAATAATACAGATAATCAACAATTAGTAGTAGCATTT<br>CTAAATGGTCTACTAGCAGGAATATGTCATTAAATTTTCTAACAATCTATTAAA<br>TTATTGTG |
| | ***GB_8.2*** | 400 | AATAATACAGATAATCAACAATTAGTAGTAGCATTTCTAAATGGTCTACTAGCA<br>GGAATATGTCATTAAATTTTCTAACAATCTATTAAATTATTGTGATTCTCTAAGA<br>TATTTGTGTTCCACTTATTTTATTAAAAACTAATATTTTTACTTGGAAGTATTTTC<br>TTTTGCTATTTCTTAACTCAGATATACCATTCCCTACGACTTGTGAAATATTTTA<br>CTAATACTTAACTAAAACTATACAAGATTTTTTCCCACTCAAGTTCTAATCAATG<br>CTTACTTTAAACACGTTTTTTTACACCTATATATACTTTAATATTTGTTTTCAATA<br>TTTTTATTTATTTGCATGCAAATTATATATCTAAAAATGGCGCCACAATGGCTGG<br>TATATAATATCCAG |
| | ***GB_8.3*** | 331 | ATTTATTTGCATGCAAATTATATATCTAAAAATGGCGCCACAATGGCTGGTATA<br>TAATATCCAGCCTGCGAGCATGCTTCAGATTTAAAGGAAAAAGGGAGAAACAG<br>CACAAAATTTGTACTTGTTAGAATCGCAAATAAATTTTGTTAAAAAAATTTGAT<br>AAAGTAGAAATTGGTGAAAAATCTCATTTTATTTGTTTCAAAAGAAAAAAATAA<br>CTTCGTATAGCATACATTATACGAAGTTATATGGTTAGCAAAGGAGAAGAGGA<br>TAACATGGCAATTATAAAAGAATTCATGAGATTTAAGGTTCATATGGAAGGTTC<br>TGTAAATGG |

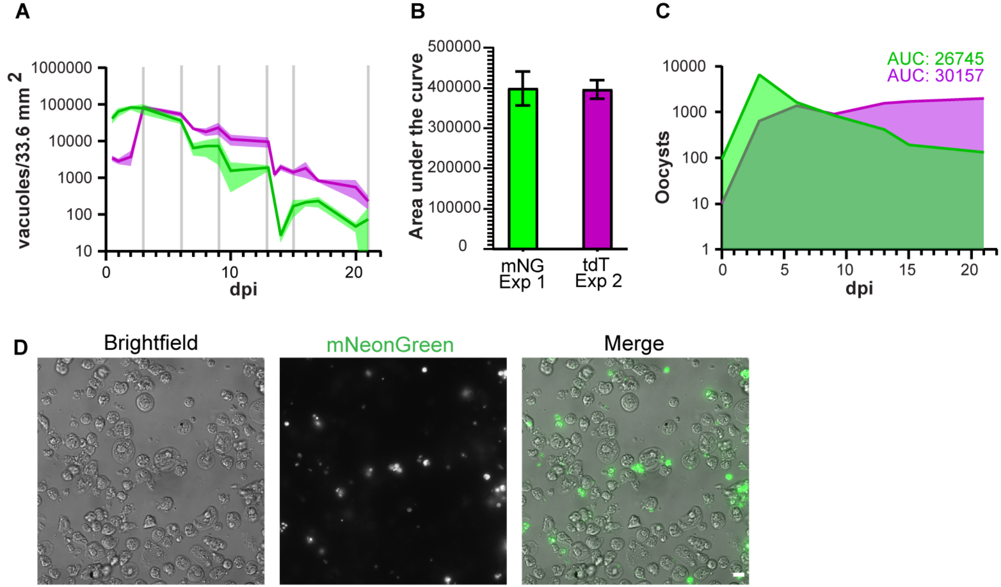


**Supplementary Figure 1 ODM infection supports long term infection and oocyst production**.(**A**) Vacuole counts from image J macro analysis of fluorescent parasite vacuoles adjusted for total cell culture area. Each colored line represents a different experiment. mNG infection (green) and tdT infection (magenta). Vertical grey lines show days with supernatant collection; images were collected before supernatants were collected. Standard deviation is shown as shaded area surrounding each line. (**B**) Area under the curve (AUC) of experiments in A. (**C**) Flow cytometry quantification of oocysts collected from ODM supernatant based on FSC and SSC of purified WT stock oocysts and expression of mNG or tdT. (**D**) Images of supernatant collected from ODMs infected with mNeonGreen expressing parasites 6 dpi. Dead host cells that have sloughed off the monolayer can be observed as well as various parasite stages (green). Parasites are observed to still be attached to host cells in many cases. Scale bar is 5 µm.

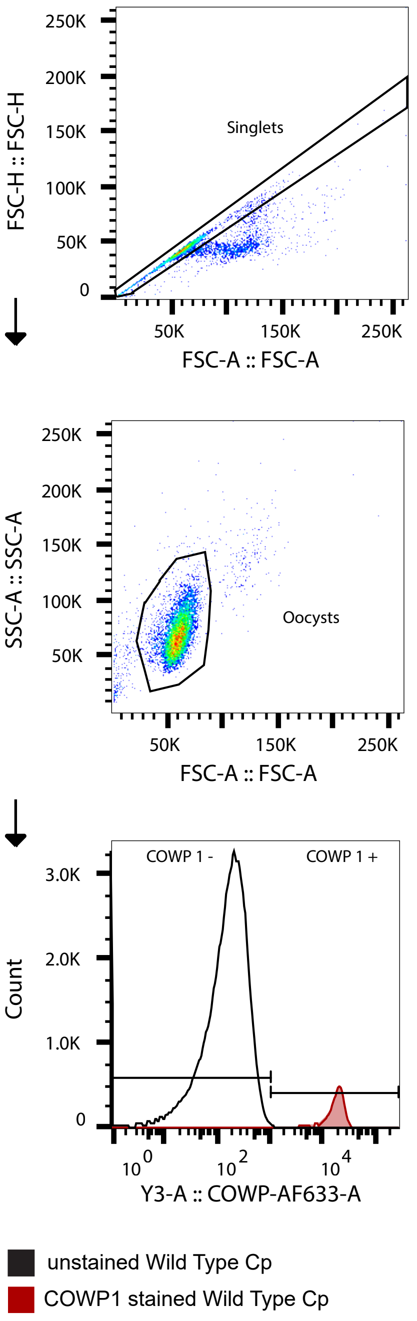


**Supplemental Figure 2. Gating strategy for oocysts used for data in figure 3.** Doublet discrimination was performed, then FSC and SSC were used to find oocyst size and granularity based on WT stock oocysts. Then COWP1 expression was used to confirm cells were oocysts. This figure was prepared with FlowJo and Adobe Illustrator.

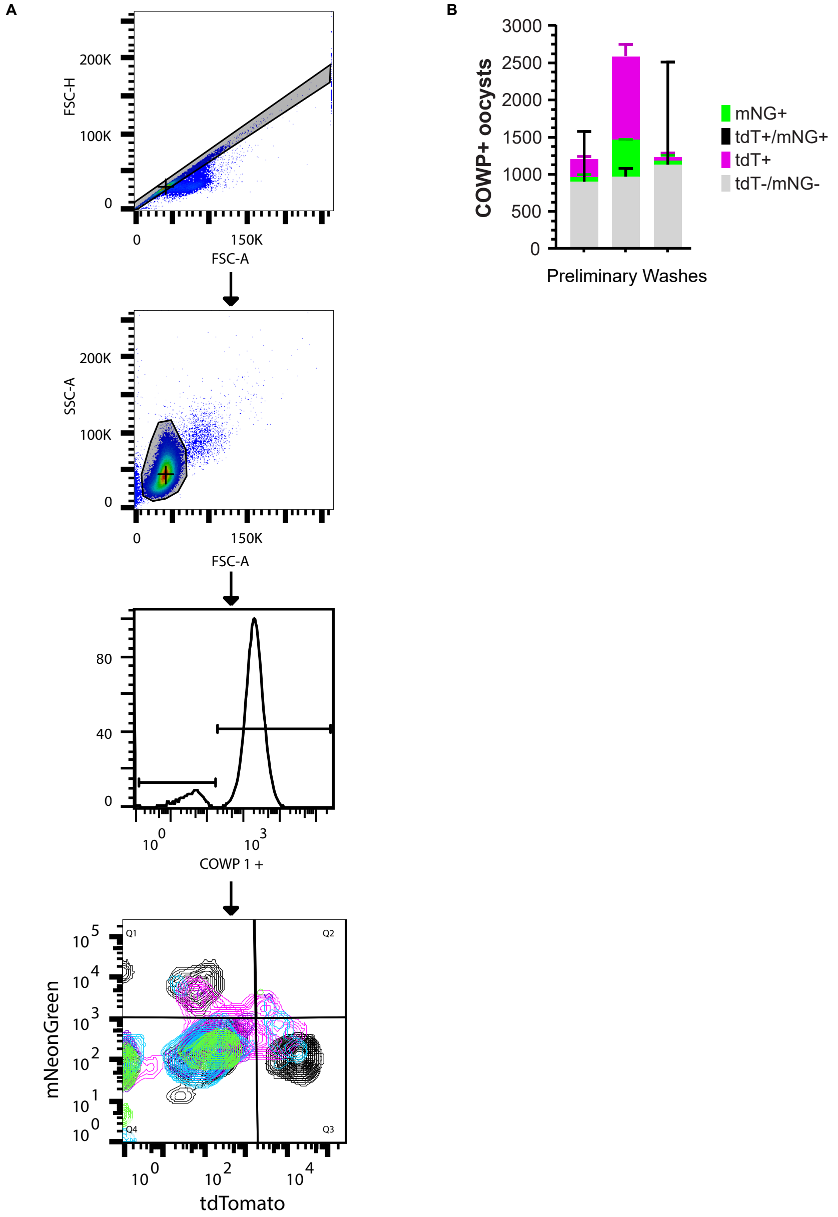


**Supplemental Figure 3. Gating strategy for mating experiments.** (**A**) Gating strategy used to measure oocyst populations in figure 4. Doublet discrimination followed by gating on FSC and SSC based on purified stock WT oocysts, then COWP1 expression to identify oocysts. To assess the subpopulations of oocysts mNeonGreen and tdTomato expression was used to identify single positive, double positive, and double negative populations. Contour plots of samples with mixed populations are shown in different colors in the last panel. (**B**) Oocyst counts from the washes done 3 hpi to remove unexcysted oocysts and unattached parasites. Washes show that either the tdT or mNG oocyst stock was either contaminated with WT or reverted to WT at some point. The mixed population of non-fluorescent and mNG oocysts in the mNG oocyst stock was visually confirmed by looking at the stock microscopically. Visual inspection of the tdTomato stock showed a pure population. This explains why the majority of the PVs counted by ImageJ macro in Figure 4 were tdT positive and that the macro counts significantly underestimate the PV count in the ODM. There was also likely crossing between WT and mNG or tdT that could not be distinguished from tdT x tdT or mNG x mNG crossing in this assay.

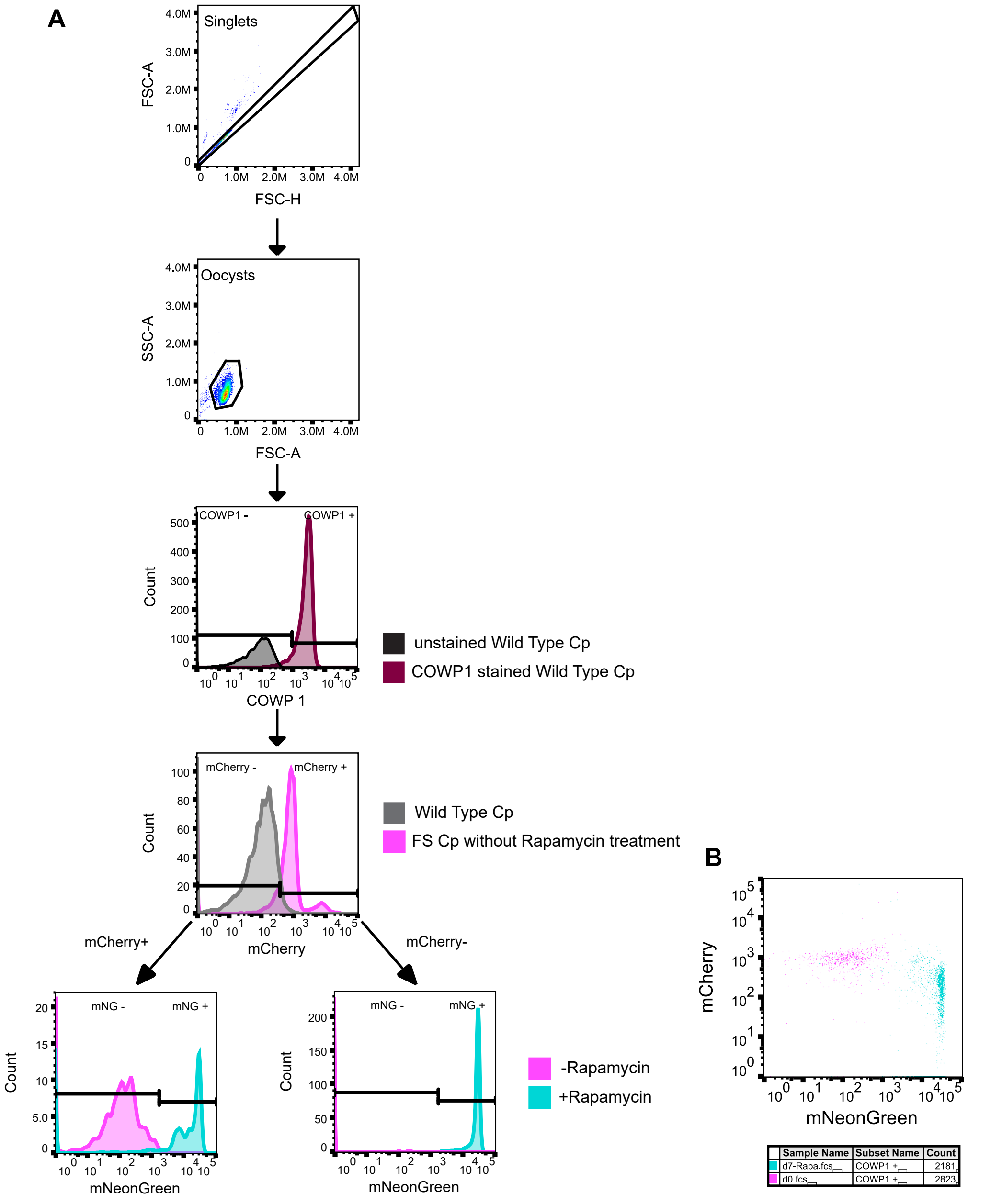


**Supplemental Figure 4. FS Oocyst gating strategy.** (**A**) Doublet discrimination followed by FSC/SSC gating based on purified stock oocysts. Then COWP+ populations were gated based on expression of mCherry or mNeonGreen. The final two histogram plots show oocysts from the untreated mouse (pink) and the treated mice (teal). mNeonGreen gates were set based on nonfluorescent stock oocysts being negative. (**B**) Oocysts from untreated mice are shown with pink dots and oocysts from mice that received rapamycin are shown in teal dots. mCherry expression is on the y-axis and mNeonGreen is on the x-axis.
